# Engineered 3D environments reprogram fibroblast-mediated inflammation in rheumatoid Arthritis

**DOI:** 10.1101/2022.12.21.521283

**Authors:** Aneesah Khan, Piaopiao Pan, Yanling Lan, Çağlar Çil, Julia Isakova, Yilin Wang, Theodora Rogkoti, Thanutchaporn Sartyoungkul, James FC Windmill, Jonathan William, Manuel Salmeron-Sanchez, Margaret H. Harnett, Oana Dobre, Miguel A. Pineda

## Abstract

Inflammation is essential for fighting infections and initiating tissue repair, but chronic unresolved inflammation underlies many conditions, like cancer and autoimmune disorders. While dysregulated immune responses drive chronic inflammation, non-immune stromal cells such as fibroblasts also play a critical role. Targeting fibroblasts could enable tissue-specific therapies while avoiding the systemic suppression caused by current drugs. However, traditional culture systems often induce artificial behaviours, limiting progress.

Here, we demonstrate the importance of the mechanical properties of the 3D culture environments in fibroblast-mediated inflammation in the context of Rheumatoid Arthritis, a chronic disease that primarily affects joints but also impacts other organs. We isolated fibroblasts from healthy and arthritic mouse joints and expanded them on 2D tissue culture plastic, stiff fibronectin-coated scaffolds, or soft pegylated fibronectin-based hydrogels.

Our results highlight the plasticity of fibroblasts, with microenvironmental cues driving inflammatory or regulatory phenotypes. The 3D environment offered by fibronectin-coated scaffolds restored inflammatory gene expression profiles that were lost in flat cultures, while the more physiologically relevant soft hydrogels shifted fibroblasts toward a resolving phenotype. These findings underscore the importance of the 3D environment in modulating fibroblast behaviour and establish a foundation for bioengineered systems that better model disease or guide therapeutic strategies.

## 1. Introduction

Stromal fibroblasts were typically described as tissue supportive cells, whose main function was to produce the matrix in which other cells were embedded. Today we face a different landscape, where fibroblasts are recognised as inflammatory mediators thanks to their ability to modulate local immune cells. Synovial fibroblasts (SFs) are a specialized population found only in the synovium, the tissue lining the inside of the capsule wrapping articular joints such as the shoulder, wrist, knee or ankle. SFs provide nutritional support and lubricating molecules like hyaluronic acid and lubricin, but they can also secrete inflammatory mediators and control bone remodelling. In inflammatory Rheumatoid Arthritis (RA), this process is dysregulated, and SFs become active disease drivers. RASFs become hyperproliferative, increase production of matrix-degrading enzymes and cytokines, and promote recruitment of immune system cells [1–4]. In turn, recruited cells secrete inflammatory cytokines, such as IL-17 or TNF, consolidating the accumulation of pathogenic factors that further activate SFs. The result is the generation of self-sustained inflammatory loops, a hallmark of chronic RA.

Clinically, modulation of SFs responses would offer a clear benefit to RA patients, as it would directly target the core disease tissue to neutralize local autoimmune networks whilst avoiding the adverse effects of current immunosuppressive drugs. However, despite intensive research in the field, no drug has been approved to target SFs yet. This reflects that our understanding of SF-dependent inflammation is incomplete, and perhaps inaccurate to some extent, as it is based not only on clinical or in vivo studies, but also on cells expanded in vitro on non-physiological two-dimensional (2D) substrates. These 2D systems have been crucial to develop the field of stromal immunology because they are easy to implement and allow ex vivo cell expansion. However, although 2D expanded SFs preserve epigenetic modifications associated with pathogenesis [5], their prolonged culture results in a gradual loss of disease phenotype [6, 7], highlighting that gene expression is adapted in response to the loss of tissue architecture and matrix-mediated signalling in the artificial flat surroundings.

In response to this, three-dimensional (3D) culture models have been engineered to mimic more physiological environments. General examples are scaffold-free derived matrices, such as collagen gelation or spheroid cell culture [8, 9]. Other techniques include protein-based structures, such as the extensively used Matrigel^®^, but also natural and synthetic hydrogels and hard polymer scaffolds [10–15]. More specific tissue-oriented approaches are 3D bioprinting designs and organ-on-chips [16–19]. The arthritic synovium represents a challenging tissue to model, due to its unique characteristics, integrating matrix remodelling and complex interactions among immune cells, the stroma and the vasculature. Early 3D SF cultures in fibrin gels were conducted over 25 years ago [20], but it is only recently that more diverse methods have been developed. Matrigel-based systems are the most widely used, supporting differentiation of specialised SF subtypes and responses to TNF [10, 21–25]. However, there are limitations in Matrigel cultures, like batch-to-batch variability and their undefined composition. More engineered approaches have included microfluidic chips incorporating fibrin gels [26], 3D printed collagen/alginate gels [27] or collagen gels [28], where the choice of the material is determined by the specific biological process to be tested. However, the global impact on cell activation and inflammatory potential is often overlooked. Furthermore, given the high degree of variability in the methods found in the literature and differences in composition in Matrigel lots, it is difficult to compare the results of these independent approaches. Thus, how differential culture conditions impact on SF cell biology outside of the primary experimental readout is unclear.

In this study, we use a murine model of arthritis to investigate the influence of ex vivo culture conditions on activated SFs. To provide a broader translational perspective, we used isolated SFs from murine synovium, both from healthy and inflamed joints, and we compared their activated phenotype ex vivo with those cells expanded in conventional 2D cultures and the same cells transferred into two distinct 3D systems: i) fibronectin-(FN)-coated polystyrene scaffolds and ii) fibronectin-(FN)-based 3D hydrogels. Scaffolds were commercially available [Alvetex^®^], and hydrogels were produced in-house, functionalizing FN with PEG-maleimide (PEGylation) as described before [29]. Our results provide a systematic comparison of the pathological signature of arthritic SFs directly isolated from the joints [not cultured, ex vivo] with those of SFs expanded in vitro in 2D culture, and following transfer to two distinct 3D systems, furthering our understanding of the effect of the culture microenvironment.

## 2. Results

### 2.1. 3D rigid scaffolds support SF culture allowing more physiological cell shape

We hypothesized that functional changes associated with SFs cultured ex vivo are due to changes in the microenvironment, rather than the cells themselves. To address this, we worked with a murine model, where SFs from mouse synovium were expanded in conventional 2D cultures [30]. Following 3 passages, some cells were transferred to 3D systems to determine microenvironmental effects. SFs expanded in 2D expressed the stromal marker vimentin at passage three and showed the expected flattened morphology, where cell adhesions are restricted to x-y plane, presenting an apical-basal polarity (Figure 1A). Since morphology can influence function, signaling and cytokine synthesis [31], first we decided to use rigid fibronectin-coated polystyrene scaffolds to add three-dimensionality without changing the physical properties of the culture material (Figure 1A, B). Scaffolds are commercially available [Alvetex^®^ [32]], engineered to a thickness of 200 μm with pores around 30-50 μm, showing a ‘sponge-like’ appearance [33]. Scaffolds supported SFs attachment and proliferation, as shown by hematoxylin/eosin staining, whilst preserving expression of vimentin (Figure 1B). Reconstruction of Z-stacked confocal images of scaffold sections stained for phalloidin presented cellular 3D morphology reflecting rounder shape, both in the cell cytoplasm and nuclei (Figure 1C). Scanning electron microscopy provided high resolution images of these 3D cultures (Figure 1D), highlighting that SFs grew both on top of the scaffold, and within it (Figure 1E) and revealing thin protrusions resembling filipodia mediating interactions with the fibronectin-coated scaffold and neighboring cells (Figure 1E).

**Figure 1.**
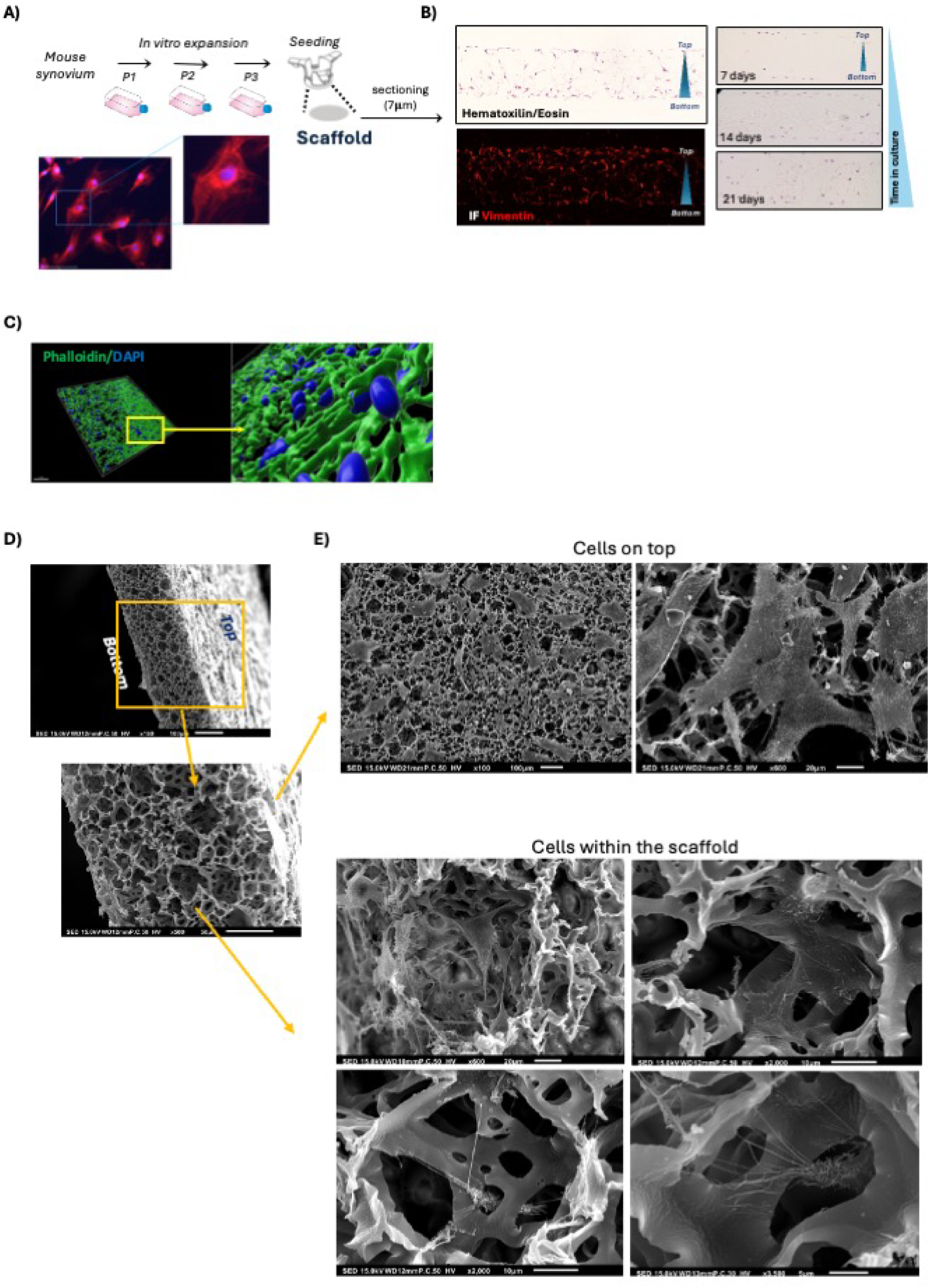
Characterization of SFs cultures in rigid scaffolds. **A)** General strategy of expansion and seeding of fibroblasts in 3D scaffolds. Images show 2D cultured SFs cells at passage three stained with anti-vimentin antibody (red) to visualize vimentin filaments. DAPI (blue) was used for counterstaining of nuclei. Scale bar: 100 μm. **B)** Hematoxylin/eosin staining and immunofluorescence to detect vimentin (red) and DAPI (blue) was conducted in sections (7 μm) of fibronectin-coated scaffolds containing naïve SFs (1×10^6^ cells, 9 days in culture). Right panel, cells in the scaffold upon 7, 14 and 21 days of initial seeding of 0.5M cells. **C)** Whole scaffolds were fixed and stained with phalloidin (green) to visualize cytoskeleton organization and cell shape prior to acquisition of Z-stack images. DAPI (blue) was used to visualize cell nuclei. Images were acquired using a Zeiss Confocal microscope (phalloidin/DAPI) staining and 3D reconstructions were done with Imaris software. **D)** Morphology of cell surface and scaffolds after 9 days in culture, assessed by scanning electron microscopy. Scale bars represent 100 μm (top image) and 50 μm (bottom image). **E)** Detailed scanning electron microscopy images of cells growing on top (Scale bars, 100 μm left panel and 20 μm right panel) and inside rigid fibronectin-coated scaffold (Scale bars ranging from 20 to 5 μm as indicated).

### 2.2 SFs culture in 3D pegylated-fibronectin hydrogels

A diverse array of gel-like systems offer an alternative to rigid scaffolds. To address modelling the joint environment on softer gels, we focused on full-length fibronectin functionalized with poly(ethylelene) glycol (PEG)-maleimide (PEGMAL). This allows covalent crosslinking of a 3D Fibronectin (FN) PEG gel with controlled chemical and physical properties [29]. Hydrogels were subsequently employed to culture cells that had been expanded in 2D (Figure 2A), employing a strategy comparable to the one previously employed for scaffolds (Figure 1A). These supported a viability of 76% after 14 days in culture (Figure 2B), with cells exhibiting morphology and vimentin expression consistent with their stromal lineage (Figure 2C). Hydrogel stiffness was measured over time, being around 4KPa for 7 days in PBS (Figure 2D). Next, we used Micro-CT to characterize the fiber and pore thickness of hydrogels (Figure 2E), which was 27.33 ± 2.22 μm and 38.04 ± 11.55 μm respectively, values resembling those reported for the 3D scaffolds. Cubic sub-volumes of interest with side lengths of 0.4 mm were selected from distinct regions (central and peripheral) within the hydrogel to capture regional heterogeneity (Figure 2F). The presence of cells in the hydrogels did not affect the porosity or the thickness of the pores, although the average number of gel fibre connections per mm^3^ (connectivity density) was increased (Figure 2G), perhaps reflecting gel degradation by secreted proteases. Overall, we demonstrated that FNPEG hydrogels support SF culture and characterised the gel properties in the presence and absence of cells.

**Figure 2.**
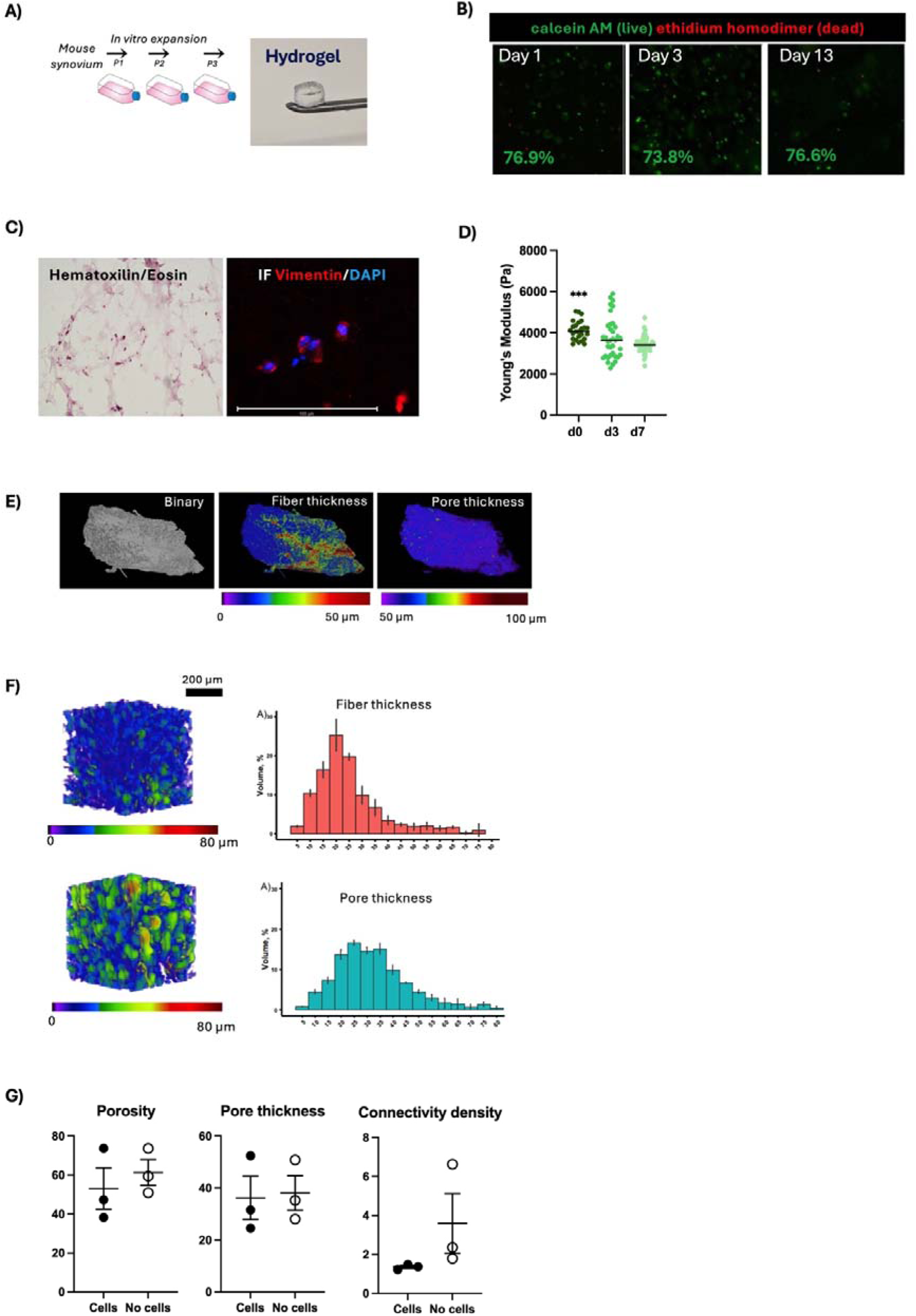
Characterization of SFs cultures in pegylated fibronectin hydrogels. **A)** General strategy of expansion and seeding of fibroblasts in hydrogels; one representative gel is shown in the picture. **B)** Cellular viability of murine SFs cultured in FNPEG hydrogels. Representative images of SFs in FNPEG gels stained with calcein AM (live, green) and ethidium homodimer (dead, red) at days 1, 3 and 7. **C)** Sections of hydrogels (7 μm) containing SFs (7 days of culture) were prepared and stained with hematoxylin and eosin and and anti-vimentin (red) and DAPI (blue) for immunofluorescence. **D)** Stiffness over time, measured by AFM nanoindentation, where x axis shows Young’s modulus (Pa) and y axis days after hydrogel formation. Error bars = SD, *** = p < 0.001 calculated by ordinary one-way ANOVA and Tukey’s test for multiple comparisons. **E)** Freeze dried hydrogels were microCT scanned to analyze morphometrical parameters. Shown: Binary image and color-coded quantified images for fiber thickness and pore size across one representative gel. **F)** Representative cubic sub-volumes of interest (sub-VOIs, 0.4 × 0.4 × 0.4 mm³) were analyzed in three independent hydrogels. Color code correlates with fiber and pore size as indicated. For histograms, x axis represents measured size in μm, y axis represents the mean of percentage of total gel volume taken for each size; error bars = SD for technical triplicates in each gel. **G)** Porosity, pore thickness and connectivity thickness were analyzed using microCT scans sub-VOIs. Each dot represents an independent gel, with (black circles) or without cells (white circles).

### 2.3. Fibroblast activation relies on environmental cues: Inflammatory signature is lost in 2D, recovered in scaffolds, and rewired in hydrogels

Once our 3D models were characterized, we compared their transcriptomic profiles of steady-state and activated SFs. To mimic human chronic inflammatory arthritis, we used the murine model of Collagen-Induced Arthritis (CIA), because it offers a well-defined and reproducible pathology to mimic the SF activation observed in human disease [34]. Our previous work has shown that SFs directly sorted from the joints of arthritic CIA mice (*ex vivo*, not cultured, to reflect their in vivo status) display increased proliferation and cytokine/chemokine activation, such as IL-17 and TNF mediated pathways. In this previous study, we identified 386 genes that are differentially expressed in arthritic SFs compared to those from healthy mice [35]. This dataset (GEO Series accession number GSE162306) provides us with a well-defined pathophysiological reference for freshly isolated arthritic SFs, being suitable for assessing the impact of subsequent in vitro expansion. We cultured SFs from both healthy and arthritic synovium in a fibronectin (FN)-coated 2D plastic until passage 3, a stage at which we have already shown SFs to retain epigenetic changes in vitro [36]. Subsequently, SFs were transferred to either FN-coated scaffolds (as in Figure 1) or FN-hydrogels (as in Figure 2) for further culture prior to transcriptomic analysis via bulk RNA-Seq. FN was chosen as the supportive matrix due to its accumulation in regions of SF proliferation in arthritic joints, where it guides SF migration to the pannus through α5β1 and αvβ3 integrin signaling and matrix metalloproteinase (MMP) action [37, 38]. Principal Component Analysis (PCA) of these RNA-Seq data confirmed that sorted arthritic SFs, ex vivo, differ significantly from their healthy counterparts, but such differences were not observed when cells were expanded in 2D (Figure 3A). Moreover, neither healthy nor arthritic 2D SFs exhibited a similar PCA profile to either sorted population (Figure 3A). Interestingly, transferring 2D expanded cells to 3D scaffolds restored the distinction between healthy and arthritic SFs. By contrast, culture in hydrogels induced distinct alterations that diverged from both the sorted and scaffold phenotypes (Figure 3A). We next evaluated the expression of genes that were differentially expressed (DE) in sorted ex vivo SFs across the rest of cultured systems, both 2D and 3D (Figure 3B). These results confirmed the loss of the ex vivo (healthy and arthritic) phenotypes during 2D expansion and that transferring SFs to 3D scaffolds restored some of differential gene expression between healthy and arthritic cells, but with responses differing in hydrogels (Figure 3B). To explore this, DE genes in arthritic SFs compared to healthy cells were evaluated in sorted ex vivo cells (Figure 3C) and 2D and 3D culture systems (Figure 3D, Supplementary tables 1-4). Key genes involved in RA progression, such as cytokines, chemokines, and genes involved in cell proliferation, were not DE in 2D expanded arthritic cells, but were re-activated in 3D scaffolds. *Ex vivo* and 3D scaffolds shared a high number of DE genes in arthritic cells, whereas most genes significantly modulated in arthritic SFs in hydrogels were unique to that group (Figure 3E). For functional studies, we conducted a comparative enrichment analysis in KEGG pathways on up-regulated genes in arthritic SFs across all culture systems (Figure 3F). As expected, SF activation was confirmed in the *ex vivo* population, showing enriched pathological pathways in two major areas, i) cytokine signaling (IL-17 and TNF) and ii) cell cycle activation and proliferation. Interestingly, these two clusters were lost in 2D cultures, but both were recovered in 3D scaffolds (Figure 3E, KEGG cluster 1). Cells in scaffolds also showed a unique group (Figure 3F, KEGG cluster 3), that was functionally related to cell proliferation. Hydrogels showed a very different scenario, where arthritic cells did not show inflammatory or proliferative signatures, but rather they displayed a significant increase in pathways related to cytoskeleton and focal adhesion reorganization, which was common to 2D, (Figure 3E, KEGG cluster 2) but also a tempered immune response (Figure 3E, KEGG cluster 4). These data reveals that the pathogenic phenotype acquired *in vivo* is heavily dependent on environmental factors, but it is not permanently lost in culture despite 2D expanded cells lose it temporally. Thus, SFs appear to be plastic, as exemplified by the recovery of inflammatory phenotype in scaffolds and the novel phenotype presented in hydrogels. An integrative analysis of Canonical Pathways, KEGG pathways and WiKi pathways supported this conclusion, showing that increased transcriptional activity of TWIST2 and decreased Jun and YBX1 responses were unique to hydrogels, whereas NFKB and STAT3 pro-arthritic responses are present ex vivo and scaffold SFs only (Supplementary Figure 1).

**Figure 3.**
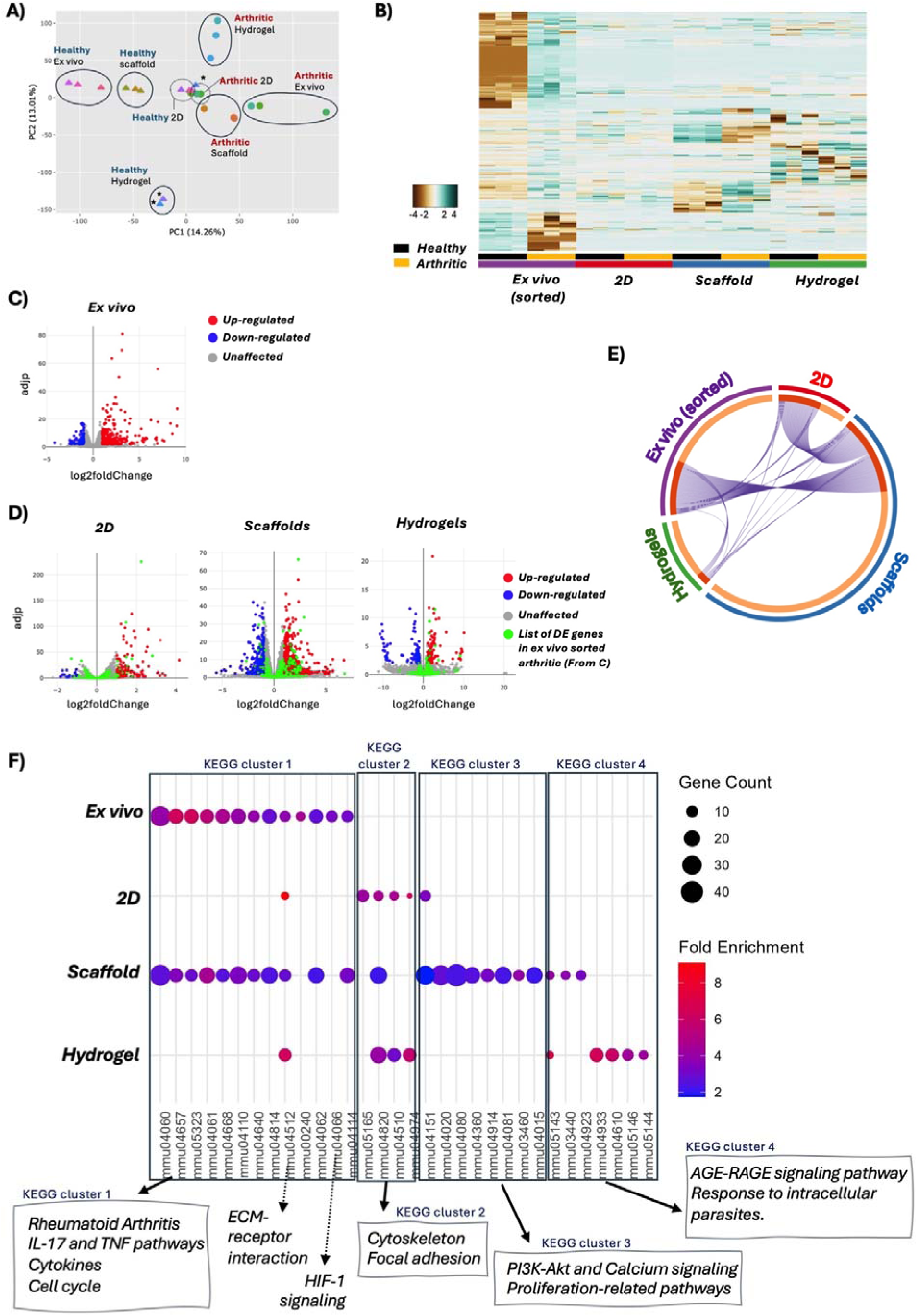
Transcriptomic changes of arthritic synovial fibroblasts across multiple culture systems reveal inflammatory memory and plasticity. SFs from healthy and arthritic mice were i) sorted by Flow Cytometry (FACs) from the joint tissue, or ii) expanded in 2D, and subsequently cultured in iii) FN-coated scaffolds and iv) FNPEG-hydrogels. RNA was isolated and analysed by bulk RNA-Seq (75 bp paired-end, 30 M reads) to compare arthritic versus naïve SFs in the four systems. Bioinformatic analysis is shown as: **A)** Principal component analysis, **B)** heatmap of expression of DE genes in arthritic cells (padj < 0.01 and |log2foldChange| > 2) in all four systems, and **C-D)** volcano plots (x = log2 Fold Change in arthritic cells, y = log10 padj value) showing significantly regulated genes (padj < 0.01 and |log2foldChange| > 2) in arthritic versus naïve conditions in cells sorted from synovial tissue (C) and expanded in vitro in 2D and 3D (D). Red dots: Up-regulated genes, blue dots: down-regulated genes. For D, green dots show the DE genes in (C) across other culture systems. **E)** Circos plot representing significantly up-regulated genes overlapping in not cultured cells (in vivo), 2D cultured cells and cells in 3D rigid scaffolds and FNPEG hydrogels. Groups represented on the arc outside; red: 2D, blue: Alvetex® scaffold, green: FNPEG hydrogels. Inner circle shows unique genes to each group in orange, and shared genes in red. **F)** Functional pathways enrichment in arthritic fibroblasts in vivo and ex vivo expanded in all culture systems. Significantly DE genes [Fold change >2, adjp < 0.05] were used, and significantly modulated KEGG pathways are shown in the x axis as pathway IDs (mmu identifiers). Circle size correlates with the number of detected genes, and the relative fold enrichment is shown in red-purple color scale as indicated in the legend. Only pathways with adjusted p values < 0.05 were included.

### 2.4. The pathogenic phenotype of arthritic SFs ex vivo is functionally recovered in vitro upon transfer to 3D scaffolds

Our RNA-Seq analysis revealed significant transcriptomic remodeling of arthritic fibroblasts in response to culture conditions. Arthritic cells isolated ex vivo showed upregulation of gene families that are fundamental for cell division [Cnn: involved in microtubule organization, centrosome assembly, Cdc: control cell cycle checkpoints, Cenp: chromosome attachment to the mitotic spindle during cell division; Kif: intracellular transport, mitosis, and cytoskeletal organization], all of them connected to Ki67-mediated activity (Figure 4A). This signature was absent in 2D and hydrogels but partially recovered in scaffolds (Figure 4B). To determine whether these changes translated to functional differences, we examined the expression of the proliferation marker KI67 (*Mki67*) at the protein level. KI67 is absent in quiescent (G0) cells but expressed in all active phases of the cell cycle (G1, S, G2, and M) to direct cell mitosis [39]. Arthritic SFs expanded in 2D did not show any significant difference in KI67 expression compared to healthy cells (Figure 4C). However, approximately 90% of arthritic SFs became ki67+ in 3D scaffolds, compared to 45% in cells from healthy animals, confirming the recovery of the proliferating phenotype in 3D scaffolds (Figure 4D). Higher KI67 expression levels were also observed, although this did not reach statistical significance (Figure 4D).

**Figure 4.**
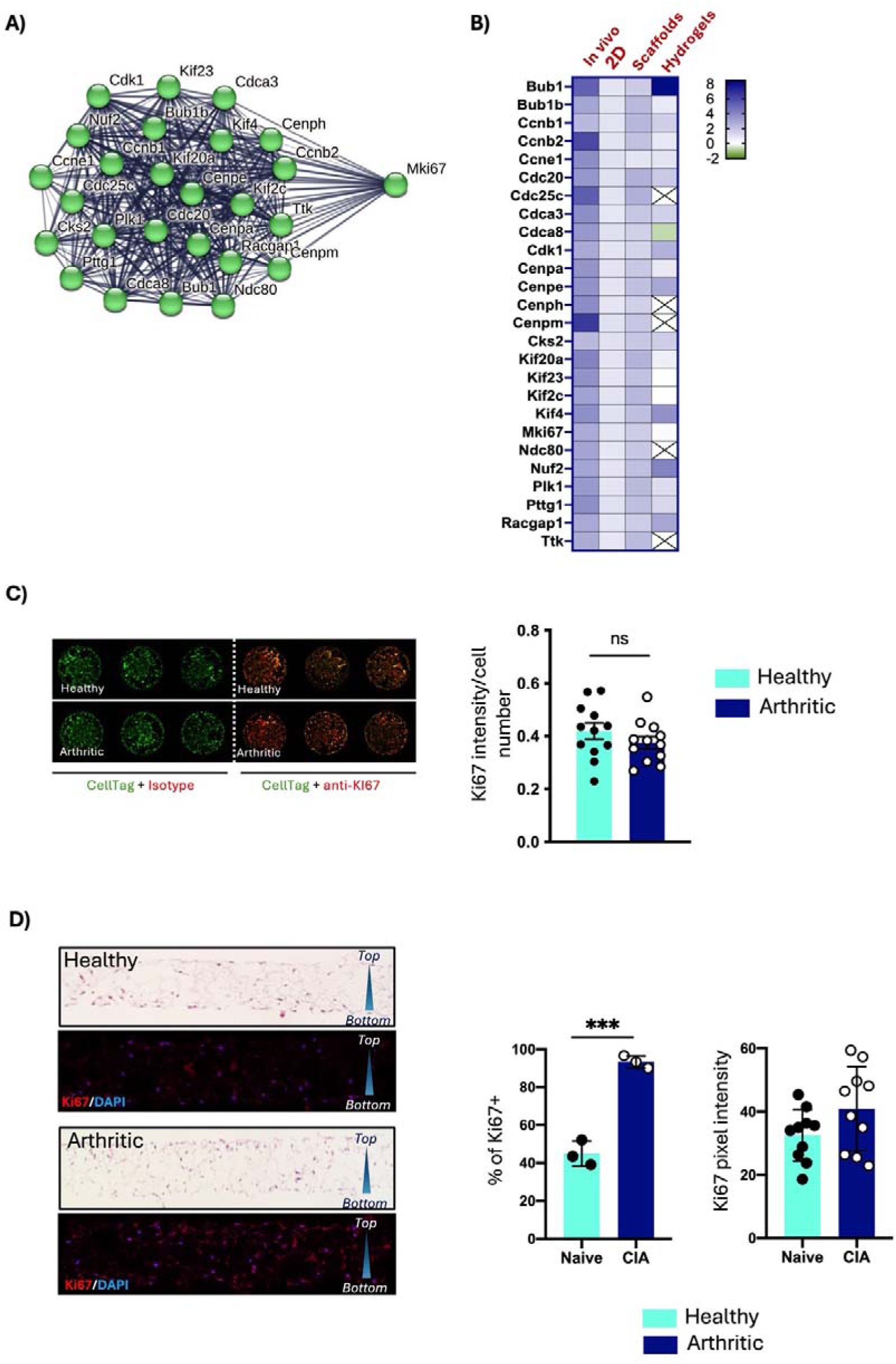
Arthritic SFs recovered the proliferating status in 3D rigid scaffolds. **A)** Significantly up-regulated genes in arthritic SFs in vivo are functionally related to Ki67 proliferation marker. Green circles represent individual genes, and lines represent functional connection identified in the literature, using STRING. **B)** Relative modulation of all genes shown in A in arthritic SFs in vivo and individual culture platforms, analyzed from RNA-Seq datasets. Results shown fold increase expression in the arthritic group compared to healthy cells. **C)** Naïve and arthritic SFs were expanded in 2D cultures. Expression of Ki67 was quantified by immunofluorescence, and signal was quantified using InCell Western. Images show a representative experiment with 3 wells. For quantification, Ki67 intensity (Red) was normalized with the number of cells (Green) in each well, represented in the column graph. Each dot represents one individual well from two independent experiments. Error bars show SD. **D)** Naïve and arthritic SFs expanded in vitro were cultured in FN-coated scaffolds for 7 days. Sections were stained with hematoxilin/eosin and antibodies against Ki67 (red) and DAPI (blue) for immunofluorescence. Percentage of Ki67+ cells and intensity of Ki67 expression was calculated and plotted in column graphs; dots represent mean of technical triplicates from three independent experiments; error bars = SD, *** = p < 0.001 calculated by two-tailed unpaired t test. Pixel intensity for Ki67 staining was calculated using image J.

While KI-67 is not a direct marker of inflammation, it is often expressed in proliferating cells within inflamed tissues. In arthritis, IL-6 and matrix metalloproteinase-3 (MMP3) are well-documented examples of inflammatory cytokine secretion and tissue damage effector pathways, respectively. In our RNA-Seq data, 3D scaffolds retrieved pathogenic expression of *Il-6* in arthritic fibroblasts and retained *Mmp3* expression, whereas hydrogels showed no activation (Figure 5A). To validate these findings, we measured secreted IL-6 and MMP3 in cell cultures (Figure 5B). Arthritic cells still showed a significant increased IL-6 expression compared to healthy controls in 2D (1.5-fold increase), but this difference was significantly higher in scaffolds (3.5-fold increase). In agreement with the transcriptomic data, arthritic cells in hydrogels not only did not up-regulate IL-6 production but rather, showed reduced expression (0.8-fold). Both 2D and scaffold systems showed elevated MMP3 expression, with no differences found in hydrogels between healthy and arthritic cells.

**Figure 5.**
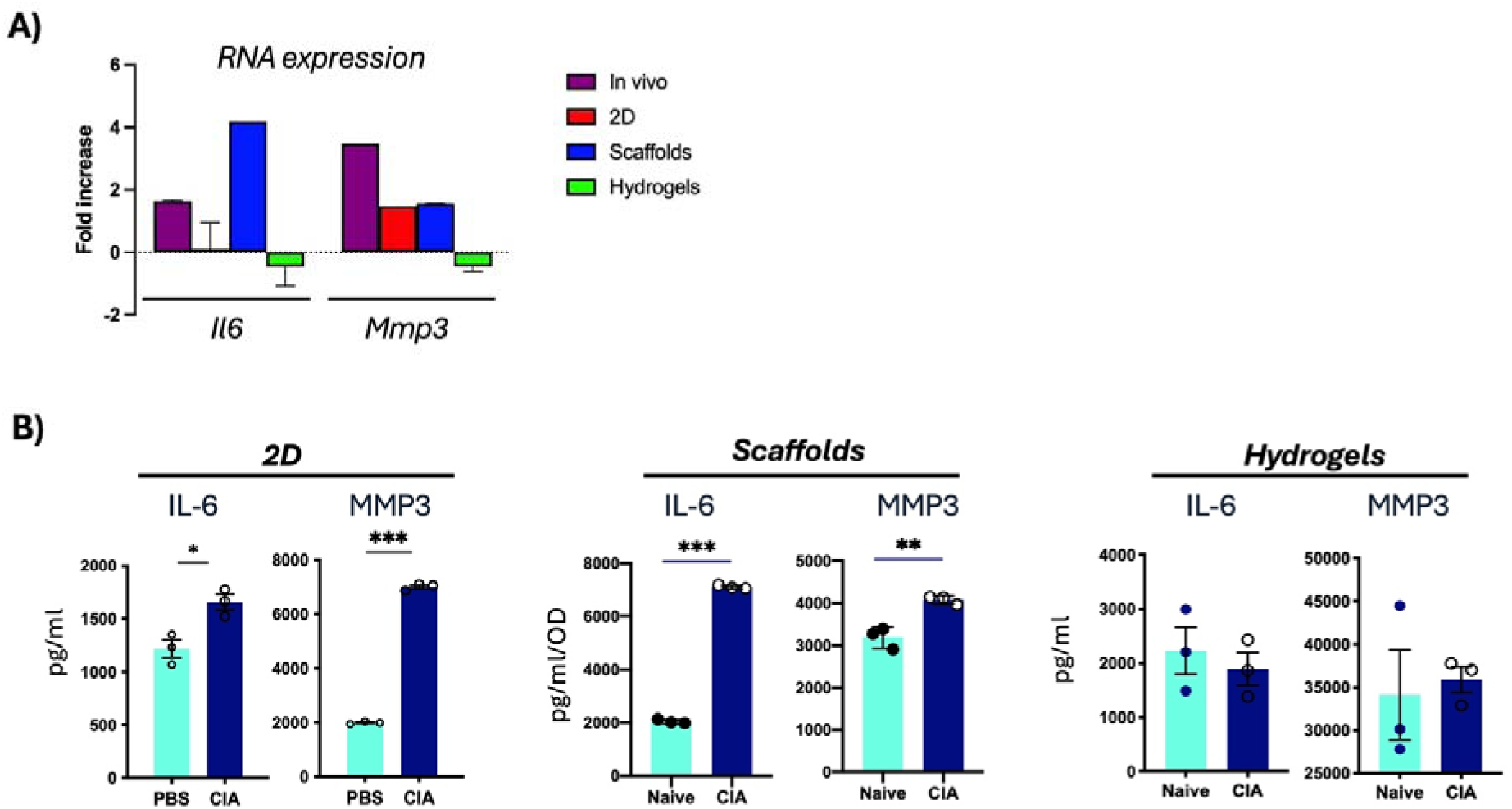
Secretion of inflammatory IL-6 and MMP3 in supernatants of cultured fibroblasts. **A)** Relative expression of *il6* and *mmp3* genes, analyzed from RNA-Seq datasets. Results shown fold increase expression in the arthritic group compared to healthy cells. **B)** Naïve and CIA SFs were cultured in 2D, FN-coated scaffolds, and non-degradable hydrogels for 7 days, when medium was replaced and collected 24 hours later to measure IL-6 and MMP3 in supernatants by ELISA. Crystal violet was used (OD 495 nm) as an indirect measure of cell number for data normalization in scaffolds. Data are presented as mean ± SD, n=3 independent experiments. Two-tail unpaired t-test was used for statistics, **p < 0.01, ***p < 0.001.

### 2.5. Fibroblasts specialization is lost in 2D culture and reinstated in 3D methods

Distinct SFs subsets are found in specific anatomical areas of the joint synovium, namely the lining and sublining membrane, and can identified by CD90^-^VCAM^+^ and CD90^+^VCAM^-^ phenotypes respectively [40–42]. SFs subsets exert non-overlapping functions, with lining fibroblasts directing tissue destruction and sub-lining fibroblasts driving inflammatory responses in RA [43, 44]. Therefore, it was key to evaluate whether culture conditions affect the ability of subset differentiation in in the cultured methods. SFs directly isolated from the joint (not cultured) of healthy mice were mostly sublining CD90^+^VCAM1^-^ (∼70%, Figure 6A), representative of the major population also described in humans [45]. A smaller subset (3.6%) was identified as VCAM-1^+^, an adhesion molecule enriched in lining areas and up-regulated in tissue-destructive SFs [41, 45]. VCAM-1+ cells went up to 8.1% in arthritic mice (2.25-fold increase, Figure 6A). However, when SFs were expanded in 2D, they presented a non-physiological phenotype dominated by CD90 and VCAM co-expression (∼60-70%, Figure 6B), which represents only a marginal population in vivo and hence such cultured SFs reflect an artefactual selective enrichment/rewiring. Cultured arthritic cells also increased VCAM1 expression compared to their naïve counterparts, albeit to a lesser extent (1.4-fold), suggesting some preservation of cell activation reflective of that occurring in the arthritic joint. The artefactual CD90^+^VCAM1^+^ phenotype in 2D may be due to polarized signaling upon interaction with the matrix in 2D. We evaluated CD90 and VCAM-1 expression in 3D scaffolds), observing a reduced percentage of 2D-associated CD90^+^VCAM-1^+^ co-expression (Figure 6C). Only ∼20% in naïve and 40% in CIA cells were found to be double positive for these markers, compared with 65% and 74% of cells in 2D (Figure 6B) and 2% and 6% in vivo (Figure 6A). Thus, culture on scaffolds was able to partially reverse the artefactual differentiation/selection of SF subtypes towards a more physiological distribution. Similar results were observed in hydrogels (Figure 6D), with more CD90+ cells than VCAM+ cells present in healthy cells, and an increase of VCAM/CD90 ratio, in line with the described pathogenic responses in vivo.

**Figure 6.**
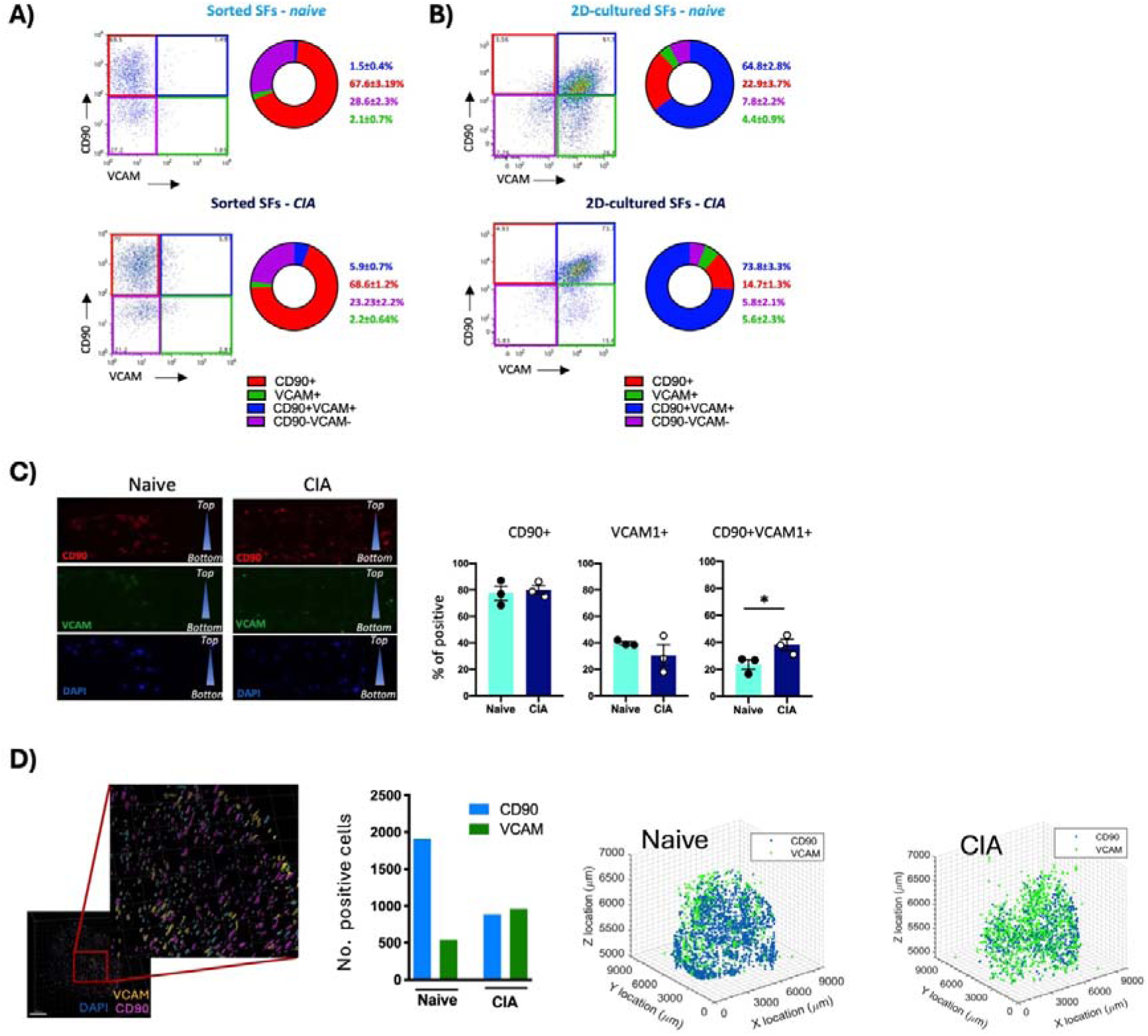
2D cultured SFs show a non-physiological phenotype reactive to fibronectin-dependent immunomodulation. **A-B)** Mouse synovium from healthy and arthritic mice was digested to obtain single cell suspensions. Expression of CD90 and VCAM-1 was evaluated by flow cytometry in joint cells prior to cultures (A) or or following expansion in 2D for 3-4 passages [2D-cultured] (B). For in vivo studies in (A), SFs were gated as live CD45-CD31-Podoplanin+. Pie charts show the proportion of CD90+VCAM-(red), CD90-VCAM+ (green), CD90+VCAM+ (blue) and CD90-VCAM-(purple) SFs. Numbers show percentage of positive SFs ± SD from 3 biological replicates. **C)** Sections of scaffolds containing SFs expanded from healthy and arthritic mice were stained for CD90 and VCAM-1 for immunofluorescence analysis. Percentage of CD90+, VCAM+ and CD90+VCAM-1+ was calculated using ImageJ. Each dot represents the percentage of positive cells shows in one independent experiment (n=3), * = p<0.05 calculated by one-tailed unpaired t test. **D)** Non-degradable FNPEG hydrogels were stained with anti-CD90 and anti-VCAM-1 antibodies, and the number of positive cells were calculated with Imaris software. Data are from one experiment, corresponding 3D scatter plots were created in Matlab using X,Y,Z positions of CD90+ and VCAM+ cells

### 2.6. SF respond to IL-1β stimulation in 2D and 3D in vitro systems

We aimed to establish whether the attenuated inflammatory phenotype observed in 2D and in hydrogels was reflective of a latent state, or whether it was the result of a general culture-dependent silencing. Therefore, we investigated whether these cultured SFs still had the capacity to respond to exogenous IL-1β, a key factor in fibroblast-mediated pathogenesis of arthritis [46, 47]. Fibroblasts responded positively to IL-1β stimulation in all cases, and the pathway ‘Rheumatoid arthritis’ was significantly enriched in SFs from each of 2D (Figure 7A), scaffold (Figure 7B) and hydrogel (Figure 7C) cultures. Other pathways enriched were related to cytokine and chemokine signalling, and JAK-STAT signalling, confirming the inflammatory activity of stimulated fibroblasts and their involvement in RA pathogenesis. Among the up-regulated genes, we detected genes related to inflammatory cytokines and chemokines *Csf3*, *Il-6*, *Ccl2* (Supplementary Table 5), shared between 2D cultures and Scaffolds (Figure 7D). Interestingly, although SFs cultured in hydrogels still responded to IL-1β stimulation, the response was significantly less robust, with less genes activated than in the other systems (Figure 7D). For example, *Csf3*, the most up-regulated gene in SFs from 2D and scaffold cultures (273-fold increase in 2D, 3816-fold in scaffolds and only 15-fold in hydrogels; Supplementary Table 5) and *Il-6* (38.2-fold increase in 2D, 528.97-fold in scaffolds and only 6.5 in hydrogels; Supplementary Table 5) highlight that whilst the scaffold SFs are the most highly responsive to inflammatory mediators, SFs from 2D culture sand to a much lesser extent, hydrogels, still retain some inflammatory capacity. The attenuated response to IL-1β in hydrogels was further confirmed by reduction in the significant enrichment pathway analysis (Figure 7E) and the absence of genes regulated by the transcription factors HIF1A, ESR1, STAT6 and SMAD3 (Figure 7F). Interestingly, IL-1β primarily activates NF-κB pathways, which are still up-regulated in hydrogels, suggesting that they could promote a broader reprogramming of fibroblasts to repress other synergistic signaling cascades.

**Figure 7:**
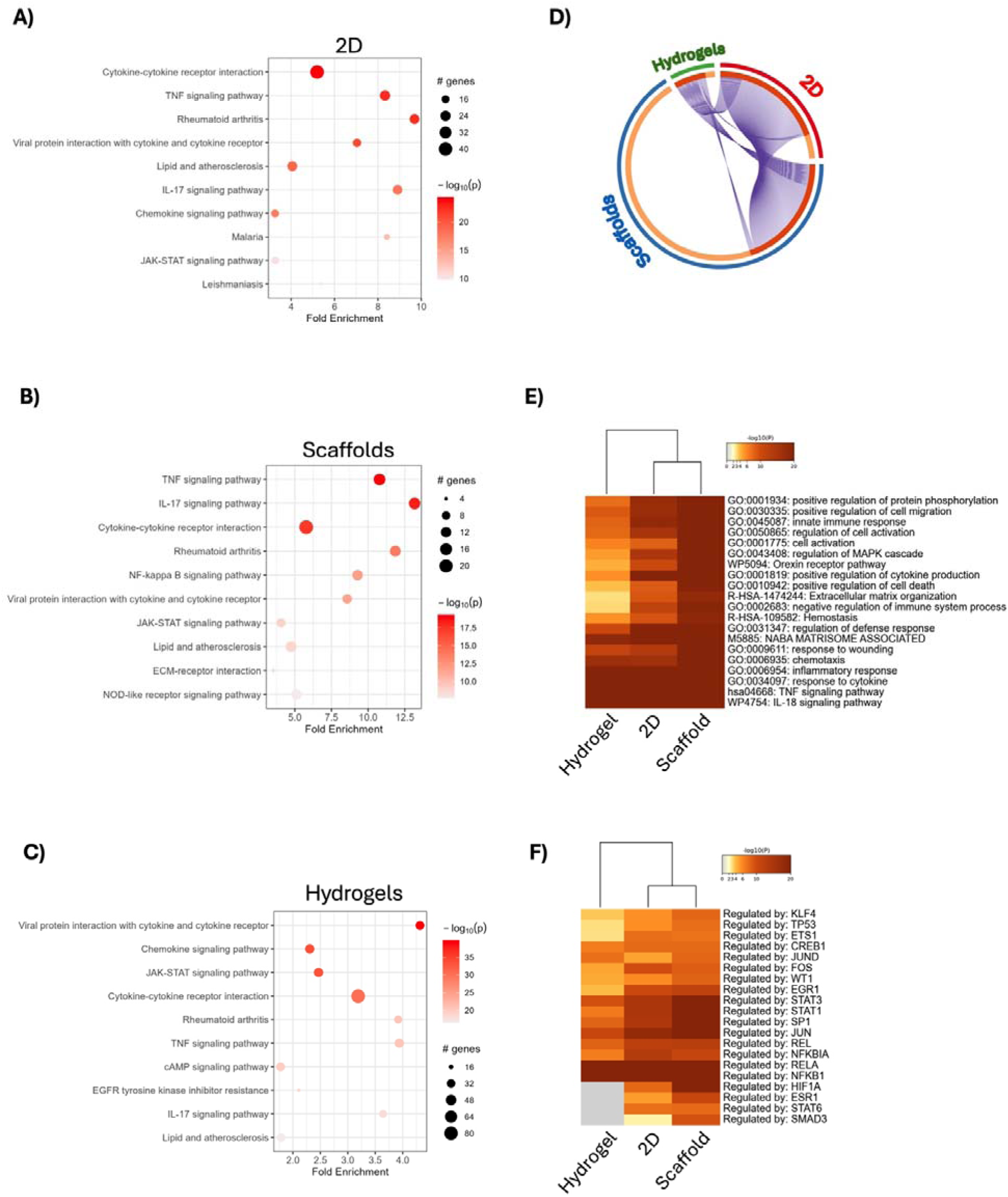
Functional pathways enrichment in healthy fibroblasts stimulated with IL-1 in 2D and 3D systems. **A-C)** Synovial fibroblasts from healthy mice were expanded ex vivo and stimulated with Il-1 for 6 hours in triplicate. RNA was then collected to conduct RNASeq transcriptomics. Significantly DE genes [Fold change >2, adjp < 0.05] were used to identify significantly modulated KEGG pathways as shown, in cells cultured in 2D (A), 3D scaffolds (B) and hydrogels (C). Circle size correlates with the number of detected genes, and the relative fold enrichment is shown in the x-axi as indicated. The –log p value is shown in red as indicated in the color scale. All pathways shown reached statistical significance (adj p <0.01). **D)** Circos plot show how IL-1 upregulated genes overlap in 2D and 3D systems (On the arc outside, red: 2D, blue: Alvetex® scaffold, green: FNPEG hydrogels). On the arc inside, each gene is assigned a spot in the arc, dark orange = genes shared in multiple lists, light orange = gene unique to that list. Purple lines link the same gene that are shared by multiple gene lists. **E)** Enriched ontology clusters in 2D and 3D systems upon IL-1 stimulation were identified and hierarchically clustered into a tree based on Kappa-statistical similarities among their gene memberships. **F)** DE expressed genes upon IL-1 stimulation were applied to TRRUST (Transcriptional Regulatory Relationships Unraveled by Sentence-based Text mining) analysis to identify activation of Transcription Factor-specific pathways based on literature curation. In heatmaps, cells are colored by their p-values; white cells indicate the lack of enrichment for that term in the corresponding gene list. All results were generated with the bioinformatic tool Metascape.

## 3. Discussion

Whilst culture on 2D resulted in a loss of functional phenotype observed ex vivo, we observed that on scaffolds, perhaps mimicking the joint/bone environment, arthritic cells partially recovered their pathogenic phenotype, indicating that this may provide a more suitable platform for candidate drug screening. Intriguingly, we found culture in hydrogels can serendipitously induce a non-inflammatory or “remission-like” phenotype that may offer new therapeutic opportunities. It is increasingly widely accepted that cells cultured in 2D change their function and biology to adapt to the new environment, including modulating the pathophysiological phenotype of inflammatory SFs [6, 7]. To date, even once recognized, the limitations of these adaptative mechanisms have been accepted in exchange for the advantages of a rapid and simple platform to study cell biology: whilst this has often led to relevant discoveries, it has likely generated some artefactual results with little or no real gain, particularly in terms of drug discovery. Thus, there is a huge effort in the scientific community to improve our current in vitro models to enhance the translational potential of basic science. However, a diverse range of methods that are not cohesive have been employed in non-comparable situations, such that it is often unclear whether cells still show physiological responses even when cultured in such more “physiologically-relevant” culture platforms. In the field of rheumatology, synovial fibroblasts play a crucial role in disease progression. However, in vitro studies have raised questions about the extent to which cellular changes are adaptations to the culture environment (2D or any 3D platform), and whether these responses are reflective of those occurring in tissues during health and disease states. We therefore chose to approach this problem using well-defined environments, where we can refer responses of cultured cells to SFs freshly isolated ex vivo to mirror the in vivo situation. Our main observation in this study is that the loss of inflammatory phenotype in arthritic SFs expanded in 2D conditions is not permanent and can be retrieved when the right environmental conditions are provided. Relocation of 2D SFs into scaffolds restored a significant number of pathogenic pathways, whilst perhaps surprisingly, FNPEG hydrogels pushed arthritic cells towards a less inflammatory phenotype. This also reveals that SFs retain a high degree of plasticity such that their phenotype is ultimately determined by ongoing signals coming from the microenvironment. Because such signals can be recapitulated by biomaterials, our findings can impact in two major areas of research, i) development of more physiological 3D culture methods to study fibroblast biology and ii) drug discovery targeting fibroblast-mediated inflammation in chronic disease.

Regarding culture methods for SFs, our findings using rigid scaffolds may lead to more physiological and affordable systems able to support pathophysiological matrix-dependent signaling and interactions. Major advances in drug discovery for RA have been made over the last decade [48, 49]. Yet, as the therapies available all target immune system cells and inflammation, they are subject to immunosuppressive side-effects that would be avoided with drugs targeting the fibroblast compartment. However, despite increasing knowledge of the role that SFs play in the perpetuation of disease, this information has not been successfully translated from the pre-clinical to clinical stage. One reason for this could be that in vitro assays for drug screening do not accurately recapitulate the synovial 3D space and associated signaling, preventing identification of relevant targets. Consistent with this, our data show that the culture conditions determine the transcriptomic and functional phenotype of SFs. The hyperproliferative and inflammatory signature of arthritic SFs is lost in 2D expanded SFs, even when epigenetic changes alter cell function and apparently stably imprint SFs [50–53]. Such epigenetic regulation includes DNA hypomethylation and microRNA expression. Indeed, the methylation pattern in RA SFs is stable over many cell passages in 2D, but these cultured cells still show a non-physiological phenotype. Interestingly, dysregulated epigenetic pathways in arthritic SFs are related to cell cycle, cell adhesion, immune function, and cytokine signaling [36]. Such pathways define a phenotype that is lost in 2D but recovered in FN-coated 3D scaffolds, suggesting that cell morphology and environmental cues and signaling are also required to establish and maintain the epigenetically imprinted inflammatory program. This could also explain why RA affect only some joints, and why DNA methylation and transcriptomic SF patterns differ between joints [54]. Establishing routine culture conditions to mimic the joint microenvironment in vitro is a priority in the field, and it must involve identification of the right structure and materials. This is especially important for samples collected at early disease stages, where the availability of biological tissue is very low and the accuracy in the diagnosis, such as disease phenotype, can be critical to efficient treatment and to prevent irreversible tissue damage. Based on our data, culture of SFs from individual patients in 3D scaffolds would allow better patient stratification and drug screening, as it could preserve the SF-dependent pathophysiology during in vitro diagnostic methods. Follow-up studies in the human context are required to validate this. Furthermore, a significant advantage of this system is its ease of implementation and relatively low-cost, which facilitates high reproducibility across laboratories, and it may represent an immediate opportunity to substantially improve conventional fibroblast culture.

Ultimately, an in vitro synovium model should consider inclusion of other cell types known to interact with SFs to create the synovial environment. Mondadori et al [26] designed a microfluidic chamber to model multi-cellular interactions, including synovial and cartilage compartments to mimic SF-chondrocyte-monocyte interactions. Similarly, Peck et al addressed the same problem, using a tri-culture platform and cell lines to model SFs and macrophages [55] whilst 3D systems to co-culture SFs and endothelial cells have been used to study cell migration and growth factors [27, 28]. Interestingly, in line with our findings, 3D environments affect the behavior of macrophage and endothelial cell lines, and synovial fibroblasts, the latter being a more aggressive RA-like phenotype [56]. These are only a few examples of recent attempts to recapitulate the complexity of joint tissues, for a more in-depth review see recent work by Johnson et al [57]. The results from these studies will be important towards the final aim of generating a synovium-on-a-chip system, although a comparative analysis of the results found in the literature may not be possible given the technical diversity employed. Currently, most research in 3D SF culture have focused on hydrogels of various nature, Matrigel being the most common choice, but also employing fibrin, collagen or alginate hydrogels. 3D hydrogels are generally used to study specific cellular responses, such as cell migration, but the effect of the culture system on global SF responses is often ignored. Our results reveal that hydrogels not only did not restore the hyperproliferative and inflammatory signature in arthritic SFs, but they rewired the already activated cells towards an altered phenotype, closer to a remission-like phenotype and distant from the original inflammatory phenotype in vivo. Matrigel has been shown to enhance expression of MMPs in RA SFs without affecting IL-6, and to allocate cells at the edge of the gels [22]. This could indicate that mechanical properties found in gel-like structures could facilitate the formation of distinct anatomical structures. Such structures have elegantly revealed novel physiological roles for Integrin α9 and Cadherin-11 [10, 22], but deeper analysis is needed to fully understand how closely gel systems can mimic synovial tissues. In this current study, arthritic SFs cultured on hydrogels showed significantly up-regulated pathways involved in osteogenesis, tissue remodeling and decreased cytokine/chemokine secretion. This difference with the rigid 3D scaffolds highlights the relevance of the culture method in SF-dependent responses. This not only should be considered in the design of 3D systems to mimic the arthritic synovium, but it also offers new avenues for therapeutic intervention, since this demonstrates that inflammatory SFs can be reprogramed. The use of hydrogels in RA models has been already tested, with specific chondroitin-based hydrogels designed to target MMP-9 production and fibroblast invasion of cartilage [58]. Although in our study we focused on FN-based environments, distinct matrisomes can be remodeled. Our RNA-Seq data showed that 2D cells lose most of the inflammatory factors, but arthritic SFs in hydrogels actively up-regulate genes not seen in 2D, nor in rigid scaffolds: for example, collagen fibrils biosynthesis and crosslinking (collagen alpha chains Col1, Col3, Col4, Col6, Col12, *P4ha*, *Adamst2*) and elastic fibre formation (Combining fibronectin, fibulins and microfibrillar-associated proteins) pathways. This could establish pro-or anti-inflammatory fibroblast-matrix crosstalk, events that contribute to RA progression at early disease stages via TLR4 signaling [59].

Moreover, the properties of materials used for tissue culture can influence FN conformation and integrin binding [60, 61] as well as functional growth factor presentation [62], which can influence cellular processes like cell migration, integrin activation and subsequent inflammatory responses. Ultimately therefore, the design of the culture system could affect/select for particular cell subset specialization, such as we have demonstrated for lining and sublining fibroblasts, that drive tissue damage and inflammation respectively [45, 63]. This hypothesis could open new bioengineering solutions, since a diversity of materials could be combined to reproduce different synovial anatomical locations in vitro. Nevertheless, the factors driving FN-dependent rewiring are still unclear and require further investigation. Differences in the stiffness of scaffolds and hydrogels suggest that this could be an important aspect, a parameter that we have not explored yet. We cannot rule out that hydrogels with higher stiffness induce a different cellular response. In fact, the arthritic joint is known to increase the stiffness of the synovium above normal values [64]. Differential mechanical forces in the joint could therefore be the required signal for SFs to start inflammatory circuits, something that could involve interactions with FN and other matrix components. Additionally, our study is restricted to fibronectin-based systems, which we have used as a model of SF enriched areas, but the synovial matrix is formed by many other components that are likely to play a role in SF-biology and disease progression. This should be considered in future studies to understand basic fibroblast pathology and to develop relevant in vitro models. Thus, recapitulation of homeostatic and inflammatory environments should be fully understood in order to develop novel in vitro systems and even therapeutic opportunities, not only in RA, but in other chronic inflammatory conditions where fibroblasts-matrix interactions play a role in disease progression. Our results highlight the importance of the culture system in the outcome of in vitro experiments, paving the way for a more rational use of these tools in rheumatology research, as well as new clinical approaches to reconfigure arthritic SFs towards a resolving anti-inflammatory phenotype.

## 4. Methods

### 4.1. Mice and Collagen-Induced Arthritis (CIA) model

Male DBA/1 mice were purchased at 8 weeks of age (Envigo; Bicester, UK) and then housed and maintained at the Central Research Facility in the University of Glasgow. All experiments were approved by and carried out in accordance with the in accordance with the U.K. Animals (Scientific Procedures) Act, 1986 and the Animal Welfare and Ethical Review Board of the University of Glasgow, UK Home Office Regulation and Licenses PPL P8C60C865 and PIL ID5D5F18C. CIA was induced using bovine Collagen type II (100 μg) emulsified with complete Freud’s adjuvant (MD Biosciences) and injected intradermally on day 0. On day 21, mice were injected intraperitoneally with a further 200 μg of collagen in PBS. Inflammation of paws was assessed every 2 days through an articular scoring system and the use of a calipers to measure paw width. Joint pathology was scored as follows: 0 = No evidence of erythema or swelling, 1 = Erythema and mild swelling confined to tarsals or ankle joints, 2 = Erythema and mild swelling extending from the ankle to the tarsals, 3 = Erythema and moderate joint swelling extending from the ankle to metatarsal joints, 4 = Erythema and severe swelling encompassing ankle, foot and digits or ankyloses of the limb. Mice were culled once stable pathology was established, typically between day 31 to day 35. A total articular index of 10 or more, paw thickness exceeded 4.5 mm or weight loss exceeded 20% of controls were considered immediate experimental endpoints.

### 4.2. In vitro expansion of SFs from murine joints

For isolation and culture of SFs, hind and front paws were harvested from mice, skin and soft tissue were removed, followed by washing 2 times in 70% ethanol and DMEM medium containing antibiotics (1% penicillin and streptavidin) before dissection. Samples were transferred paws to DMEM containing 10% FCS, 1% antibiotics, 1% nystatin, 1% glutamine and 1mg/ml type II collagenase (Sigma). 1 mg/ml DNase I (Sigma) was added for FACs experiments. Samples were incubated with constant shaking at 37°C for 80 min. After digestion, EDTA was added at a final concentration of 0.5mM and incubated at 37°C for 5 min. Following completion of the incubation time, samples were vortexed vigorously to release cells. Cells were then centrifuged, and supernatant was discarded. For cell expansion, cells were resuspended in complete DMEM medium (10% FCS, 1% penicillin and streptavidin, 1% L-glutamine and 1% non-essential aminoacids) and seeded in culture flasks. Culture medium was refreshed after 24 hours. Cells were fed twice a week and passaged when they reached 90% confluence using trypsin EDTA (ThermoFisher Scientific). Cultured cells were used at passage 3 or 4. Cultured cell purity was accessed by flow cytometry before experiments, CD11b+ cells were removed using magnetic column. Phenotype was evaluated following trypsinization by Flow Cytometry using anti-mouse podoplanin (PDNP)-alexa 647, anti-mouse CD11b-FITC, anti-mouse CD90PECy7 and anti-mouse VCAM-1-FITC antibodies. Zombie Violet dye (Biolegend) was used to exclude dead cells from analysis. All antibodies were purchased from eBioscience. For *in vitro* stimulation of SFs, recombinant IL-1β and IL-17 (immunotools) were used at 10 ng/ml for 12 hours to measure cytokine release by ELISA or 6 hours for transcriptomic experiments.

### 4.3. Culture of expanded SFs in Alvetex^®^ scaffolds

Alvetex® scaffolds (AMS Biotechnology (Europe) Lt, Abingdon) were rendered hydrophilic with 70% ethanol. The ethanol was removed, and the scaffolds washed with PBS twice (for ∼1 min) and aspirated. The scaffolds were then coated with fibronectin (RD System, 1030-FN) in PBS (0.5 mg/ml) and incubated at room temperature for 1 h in a well plate. The FN solution was then aspirated and replaced with complete DMEM medium. Scaffolds were then seeded with cells (1 x 10^6^ cells per 6-well plate scaffold), with the cells dispensed at the centre of the scaffold and then incubated at 37°C, 5% CO_2_ for 2 h to facilitate cell attachment. The cells were then gently flooded with cDMEM medium and cells were then grown on the scaffold for the indicated days, with the medium being replaced every second day.

### 4.4. Preparation of FNPEG hydrogels

FNPEG hydrogels were prepared as previously described [29], following the subsequent steps: 1) Denaturation of FN. Fibronectin (FN) (YoProteins, 3 mg mL^−1^; 50 μg per 50 μl hydrogel) was pegylated. FN was denatured in denaturing buffer (5 mM Tris(2-carboxyethyl)phosphine hydrochloride [TCEP, pH 7, Sigma] and 8 M urea [Fisher, 18 M stock] in phosphate buffer saline [PBS, Gibco, pH 7.4]) for 15 min at room temperature. 2) Protein PEGylation. An appropriate amount of 4-arm-PEG-Maleimide (PEGMAL, LaysanBio) was incubated for 30 min at room temperature at a molar ratio FN:PEGMAL 1:4. The tubes were then placed on a rotating platform (100 rpm) at room temperature for 30 min. The reaction was stopped with 2 μl NaOH at 1M. 3-Protein alkylation. The remaining non-reacted cysteine residues were blocked by alkylation using 14 mM iodoacetamide (IAA, Sigma) in PBS at pH 8 (on a rotating platform (100 rpm)) for 2 h at room temperature. The product of the reaction was dialysed (Mini-A-Lyzer, MWCO 10 KDa, ThermoFisher) against PBS for 1 h at room temperature. 4-Protein precipitation. Nine volumes of cold absolute ethanol were added to the protein solution. The mixture was then incubated at −20°C overnight and centrifuged at 15,000 g and 4°C for 15 min. The supernatant was discarded, and the protein pellet was further washed with 90% cold ethanol and centrifuged again at 15,000 g and 4°C for 5 min. Pellets were dried and solubilized using 8 M urea at a final protein concentration of 2.5 mg/mL. Once the protein was dissolved, the solution was dialysed against PBS for 1 h and stored in the freezer. 5-Hydrogel formation. PEG hydrogels were formed using the Michael-type addition reaction under physiological pH and temperature. Briefly, a final concentration of 1 mg/mL of PEGylated FN was added to PEGMAL. Following this, the thiolated crosslinker was added, at a molar ratio 1:1 maleimide:thiol to ensure full crosslinking. The crosslinkers used were either PEG-dithiol (PEGSH, 2 kDa, Creative PEGWorks; for non-degradable gels) or mixtures of PEGSH and protease-degradable peptide (VPM peptide, GCRD<u>VPM</u>SMRGGDRCG, purity 96.9%, Mw 1696.96 Da, GenScript; for degradable gels). Degradable gels were used for experiments requiring RNA extraction. SFs were mixed with the protein and PEGMAL before addition of the crosslinker. To allow gelation, samples were incubated for 1 h at 37 °C once the crosslinker was added. Hydrogels were 5% FNPEG with 0.5% VPM for degradable gels.

### 4.5. Microcomputed Tomography (μCT)

For hydrogel preparation, hydrogels (n=3 hydrogels per group) were fixed in formalin and left to equilibrate in PBS for 24 hours at 37°C. The following day, hydrogels were snap-frozen in liquid nitrogen and left to lyophilise for 24 hours prior to scanning. The lyophilised hydrogels were μCT scanned (Skyscan 1172, Bruker) with a voxel side length of 2.5µm. Scans were performed at 42 kVp X-ray tube voltage, 100 µA tube current, 1000 ms exposure time, 0.3° rotation step (for 180° total), without a metal filter and with frame averaging of two. Scans were reconstructed into grayscale images using Skyscan NRecon software (Bruker, Version 1.6.9.18). For each sample, three representative cubic sub-volumes of interest (SubVOI) with side lengths 0.4 mm, were selected for 3D morphometric analysis in CT-Analyser software (Version 1.20.8). SubVOIs were binarised using automatic Otsu thresholding (in 3D space) and denoised (removal of black speckles smaller than 25 voxels and white speckles less than 50 voxels). The morphometric parameters porosity (%), average gel fibre and pore thickness (µm) (and their distributions calculated using the maximum sphere fitting algorithm (Hildebrand & Rüegsegger, 1997)), and connectivity density (mm-3) were calculated in CT-Analyser.

### 4.6. RNA extraction and RNA-Seq

RNA from cells in 2D and scaffold cultures were extracted using RNeasy MiniElute spin columns (Qiagen) according to manufacture’s instructions at the indicated times. RNA was extracted from degradable hydrogels containing VPM and lacking PEGSH. After 3 days in culture, the hydrogels were removed from their inserts and placed in tubes with an equal volume of 2.5 mg/ml collagenase in PBS and incubated at 37°C for 90 min, gently pipetting samples slowly every 30 min to help release cells. Following incubation, the samples were pipetted vigorously and passed through 100 μm cell strainers into tubes and then centrifuged at 500 g for 10 min at 4°C. RNA was extracted using Qiagen kit, as per manufacture instructions. RNA integrity was checked using the Agilent 2100 Bioanalyzer System. Samples with a RIN value >9 were used for further experiments. For 2D and Alvetex® samples the libraries were prepared using the TruSeq mRNA stranded library preparation method. For FNPEG hydrogels the libraries were prepared using the NEBNext Single Cell-Low Input RNA Library Prep Kit for Illumina. All samples were sequenced 2[×[75[bp to an average of more than 30 million reads. RNA-Seq reads were then aligned to the mouse reference genome (GRCM38) using Hisat2 version 2.1.0, and featurecounts version 1.4.6 was used to quantify reads counts using the Galaxy portal (University of Glasgow). Data quality control, non-expressed gene filtering, median ratio normalization (MRN) implemented in DESeq2 package, and identification of differentially expressed (DE) genes was done using the R Bioconductor project DEbrowser [65]. Genes that passed a threshold of padj < 0.01 and log2foldChange[>[2 in DE analysis were considered for further analysis. KEGG pathway enrichment analysis was performed using the enrichKEGG() function from the clusterProfiler package. Pathways were considered significantly enriched if both the adjusted p-value and q-value were below 0.05. The enrichment results from each condition were converted to data frames and annotated with their respective experimental group (e.g., “2D SFs”, “Hydrogel SFs”). Visualization was carried out using ggplot2 and enrichplot. The Cytoscape [66] plug-in ClueGO [67] was used for analysis and visualization of enriched pathways in DE genes. Gene ontology terms from KEGG pathways, Wikipathways and CORUM were used to analyze enriched pathways and group functionally related networks. Metascape software [68] was used for multi-system comparisons in the generation of circos plots and identification of statistically enriched terms in GO/KEGG terms, wikipathways, GSRA and GO_TRRUST.

### 4.7. Immunohistochemistry of scaffolds and hydrogels

For preparation of scaffold blocks and sections, culture medium was aspirated after culture at indicated times and washed 3 times with cold PBS. The scaffolds were then removed from their inserts and fixed with 4% PFA at 4°C for 12h. The fixative was aspirated, and the scaffolds were washed with PBS and dehydrated using a series 30%, 50%, 70%, 80%, 90% and 95% solutions of ethanol, 15 min each, and 100% absolute ethanol for 30 minutes. Scaffolds were embedded using a Lecia Asap 300 processor (Histoclear [Merk] 30 min, 50:50 solution of Histoclear:paraffin wax 50:50 ratio 60 °C for 30 min and Histoclear:paraffin wax mix 60°C for 60 min). The wax was left to cool and set at room temperature for 2 hours. Sections were cut at 7 μm thickness for further analysis. Hydrogels were fixed in 4% PFA for 30 min, following Hydrogels dehydration in 30% sucrose solution in PBS overnight at 4°C and subsequently embed in Tissue-Tek^®^ O.C.T. compound (Sakura). Samples were placed immediately at -80°C overnight prior to cutting sections at 20 μm.

### 4.8. Immunofluorescence

For cell staining, SFs (2000-5000) were seeded on a chamber slide and fixed with 4% PFA for 10 min at room temperature. Cells were washed three times with PBS and permeabilized with PBS 0.05% Triton X-100 for 10 minutes and blocked (PBS 1% BSA) for 1 hour. Slides were washed three times in PBS 0.05% Tween 20 (PBS-T) before being incubated with primary antibody overnight at 4°C. Primary antibodies used were: anti-vimentin (Sigma); anti mouse CD90 (Biolegend), anti-mouse VCAM-1 (eBioscience) and anti-mouse Ki67 (Abcam). Slides were then washed three times in PBS-T and incubated with fluorochrome conjugated secondary antibodies for 2 hours at room temperature. Phalloidin-FITC (Thermofisher Scientific) was used to visualise actin. Slides were mounted with SlowFade™ Diamond Antifade Mountant with DAPI (ThermoFisher Scientific) to stain nuclei and covered with a glass cover slip. Slides were visualized with an EVOS^TM^ FL Auto 2 microscope. For imaging of whole hydrogels, gels were fixed with 4% PFA for 30 min at room temperature. Permeabilisation of the cells was carried out for 15 min at room temperature using 0.1% Triton in PBS, followed by washing once with PBS. The hydrogels were then treated with PBS 1% BSA for 30-60 min at room temperature, after which the primary antibody was added and left overnight at 4°C. Samples were treated with the appropriate secondary antibody for 1 h at room temperature and washed with PBS 0.5% Tween20 before mounting onto a glass bottom petri dishes using Vectashield slow fade media containing DAPI (VectorLabs). Scanning and imaging was carried out on a Leica DM8 widefield microscope using LAS X Life Science software at a magnification of x10. The hydrogels were imaged in a tilescan format with z-steps of ∼10 μm to obtain stacked images, which were then merged. The image was then deconvoluted using Huygens essential after which it was processed using IMARIS (“Cell biologist package”) to obtain 3D reconstructions and spacial cell analysis.

### 4.9. Scanning electron microscopy (SEM)

Alvetex® scaffolds were rendered hydrophilic using 70% ethanol, followed by two PBS washes and aspiration. The scaffolds were then coated with fibronectin (YoProtein, 633) in PBS (0.5 mg/mL, 300 μL per disc) and incubated at room temperature for 1 hour. After aspiration, the coating solution was replaced with complete Dulbecco’s Modified Eagle Medium (cDMEM). Cells (1 × 10[ per 12-well plate scaffold, 70 μl) were seeded at the scaffold center and incubated at 37°C, 5% CO[ for 2 hours to facilitate attachment. The scaffolds were then flooded with cDMEM and cultured for the indicated duration, with media changes every other day. For SEM, cultures were washed twice with PBS, and scaffold sections (2–3 mm²) were excised. Samples were fixed in Karnovsky’s fixative (4°C, 90 min), washed with 0.1 M phosphate buffer, and post-fixed in 1% osmium tetroxide (4°C, 90 min). Dehydration was performed using graded ethanol (30% to absolute), followed by critical point drying. Finally, samples were gold-coated via sputter coating and imaged using a TESCAN CLARA Ultra High Resolution Scanning Electron Microscope.

### 4.10. Hematoxilin/Eosin staining

For paraffin sections, slides were heated in an oven at 60°C for 35 min to melt wax. Sections were immersed in xylene to dewax and in graded ethanol solution (100%, 90% and 70%) to hydrated. Then washed in running water to remove all reagents. Sections were then stained in Harris Haematoxylin for 2-3 min and washed under running water to remove excess dyes. Staining background was reduced by dipping section in 1% acid for a few second, quickly rinsed in running water, immersed in Scott’s Tap Water Substitute for 30 seconds and then quickly rinsed in running water. Counter staining was then carried out by dipping sections in 70% ethanol for 9-10 times and immersed in 1% Eosin for 2-3 min. Sections were then dehydrated followed by immersed in 90% ethanol for 1 min, 100% ethanol for 6 min and 100% xylene for 6 min. Sections were then mounted with DPX mountant and sealed with coverslips. For staining of frozen section such as hydrogels, slides were fixed in ice-cold acetone/ethanol (75%/25%) for 10 min at room temperature, then left to air dry for 10 minutes prior to staining as described for hydrated paraffin samples.

### 4.11. Enzyme-linked immunoassays (ELISA)

IL-6, CCL2 and MMP3 secretion was detected using ELISA kit (R&D, Dou set) following manufacturer’s instructions. SFs were seeded in 96 well plates (Corning) at 10,000 cells per well in 200 μl cDMEM. For scaffolds, cell number was quantified using crystal violet (Sigma). Cells were stained with 100 μl of crystal-violet (0.04 mg/ml) for 30 min at room temperature, then washed at least three times to remove any residual dye. 1% SDS solution (Sigma) was added next to dissolve cells for 1 hour in a shaker. Absorbance was read at 595 nm using a Tecan Sunrise plate reader.

### 4.12. Flow Cytometry

Cultured cells were detached from the plates by trypsin treatment. Cells were stained with Zombie Violet (Biolegend) or flexible viability dye eFluor 780 (eBioscience) at 1 μl/ml in PBS for 20 min on ice to exclude dead cells. Cells were then washed three times with FACS buffer (PBS with 0.5% FCS and 2mM EDTA), and Fc receptor was blocked using CD16/CD32 specific antibody for 20 min on ice. Primary antibodies used were anti-CD31 (Invitrogen), anti-CD45 (Biolegend), anti-CD90.2 (Biolegend), anti-VCAM-1 (Biolegend) and anti-PDPN (Biolegend). Data were acquired using a BD LSRII flow cytometer and analyzed with FlowJo software 8.7.3.

### 4.13. In-cell Western assay

Naïve and CIA murine fibroblasts from passage 3 were seeded in tissue culture treated 96 well plates (Corning) at 6,000 cells per well in 200µl and allowed to attach overnight. Cell cycles were then synchronized by serum-starving cells for 24h (1% FBS in cDMEM) and further culturing cells for 48h, until confluent. Cells were fixed with PFA for 15 mins. Fibroblasts were gently washed two times with PBS and permeabilized with PBS 0.1% Triton X-100 for 15 mins. Two further washes of PBS 0.05% Tween and one wash with PBS were done before blocking wells in PBS 1% BSA for 30 mins. Biotinylated Ki-67 primary antibody (ThermoFisher Scientific) was applied, and cells were incubated at 4°C overnight. Wells were washed with PBS before applying IRDye 800CW Streptavidin (Li-Cor) secondary antibody and Cell Tag 700 (Li-Cor) cell stain and incubating for 1h at room temperature. Plates were then washed twice with PBS and once with PBS 0.05% Tween, and left to airdry for 15 minutes, protected from light. Measurements were taken using Li-Cor Odyssey M imager and analysed with Empiria Studio® Software (Version: 3.2.0.186).

### 4.14. Statistical analysis

Data are presented as the mean ± standard deviation (SD). Statistical analysis was performed with Prism 8 software (GraphPad). One-way analysis of variance (ANOVA) was used to test for comparing differences among multi-groups, student t-test was used to show differences between two study groups. P values <0.05 were considered significant (*).

## 5. Data availability

All data created or used during this study will be openly available from public repositories. Transcriptomic data, including raw sequencing data and processed read counts for each sample will be deposited in NCBI’s Gene Expression Omnibus.

