## Supplemental tables and figures for "Engineered 3D environments reprogram fibroblast-mediated inflammation in rheumatoid Arthritis"

### Supplementary Table 1: Top 50 up-regulated genes CIA SFs (sorted in vivo)

| ID | log2FoldChange | foldChange | log10padj |
|---|---|---|---|
| Il1rn | 9.118 | 555.467 | 29.240 |
| Olfm4 | 9.077 | 540.043 | 16.097 |
| Il18rap | 8.599 | 387.820 | 10.227 |
| Reg3g | 8.388 | 335.016 | 17.556 |
| Saa3 | 7.022 | 129.987 | 18.293 |
| Cxcl5 | 7.004 | 128.367 | 57.776 |
| Csf3 | 6.966 | 125.004 | 2.743 |
| C430002N11Rik | 6.870 | 117.003 | 7.359 |
| Prokr2 | 6.282 | 77.836 | 20.536 |
| Csf3r | 5.858 | 58.002 | 6.574 |
| 5730559C18Rik | 5.227 | 37.445 | 14.359 |
| Uox | 5.005 | 32.113 | 8.893 |
| Il1b | 4.599 | 24.236 | 8.844 |
| Gm44439 | 4.418 | 21.375 | 3.084 |
| Lilr4b | 4.206 | 18.454 | 3.620 |
| Acat3 | 4.037 | 16.421 | 21.570 |
| Hp | 3.948 | 15.432 | 17.168 |
| Spink6 | 3.710 | 13.086 | 7.940 |
| Lman1l | 3.709 | 13.080 | 11.926 |
| Hdc | 3.684 | 12.850 | 4.717 |
| Ephx3 | 3.609 | 12.200 | 6.136 |
| Saa1 | 3.472 | 11.100 | 26.425 |
| Mmp3 | 3.470 | 11.081 | 19.715 |
| Fcgr2b | 3.465 | 11.044 | 3.739 |

| ID | log2FoldChange | foldChange | log10padj |
|---|---|---|---|
| Acod1 | 3.403 | 10.575 | 5.959 |
| Lcp1 | 3.392 | 10.500 | 6.834 |
| Timp1 | 3.178 | 9.051 | 83.526 |
| Col10a1 | 3.155 | 8.907 | 5.102 |
| Ptprn | 3.134 | 8.782 | 6.464 |
| Lcn2 | 3.118 | 8.681 | 71.395 |
| Saa2 | 3.065 | 8.371 | 2.717 |
| Bank1 | 3.057 | 8.322 | 2.903 |
| Rtn4rl2 | 3.050 | 8.282 | 6.317 |
| Gja4 | 3.005 | 8.029 | 4.791 |
| Apln | 2.998 | 7.991 | 28.211 |
| Kif26b | 2.995 | 7.971 | 11.711 |
| Cdca7l | 2.935 | 7.647 | 5.590 |
| Chl1 | 2.916 | 7.550 | 8.130 |
| Gm48878 | 2.913 | 7.532 | 6.464 |
| Fam110c | 2.862 | 7.271 | 2.074 |
| Hcar2 | 2.804 | 6.985 | 5.875 |
| Ptx3 | 2.796 | 6.947 | 50.792 |
| Cenpm | 2.774 | 6.839 | 2.207 |
| Gpr39 | 2.670 | 6.364 | 4.021 |
| Ccnb2 | 2.617 | 6.134 | 10.224 |
| Slc15a3 | 2.616 | 6.130 | 7.301 |
| Mogat2 | 2.606 | 6.087 | 4.559 |
| Cbr2 | 2.605 | 6.083 | 4.680 |

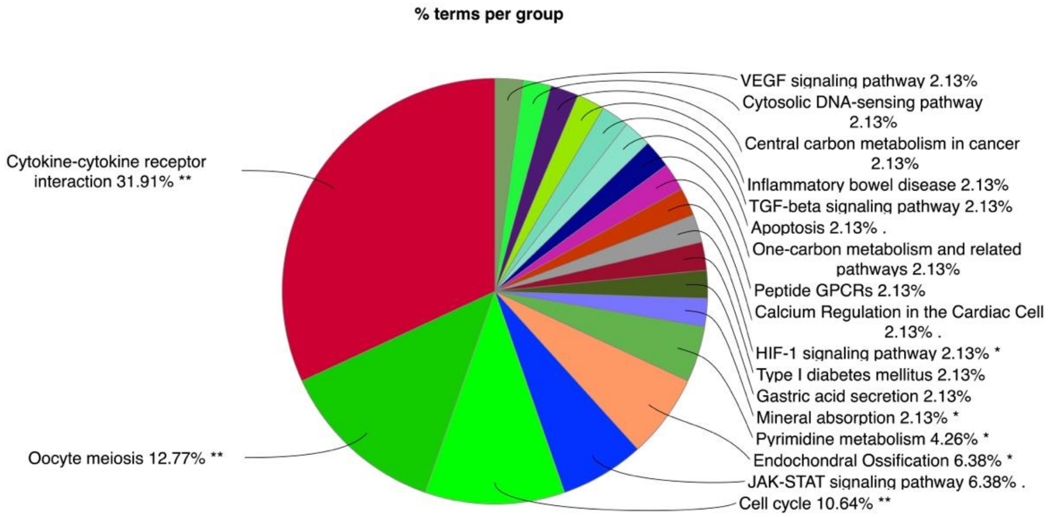

Supplementary Table 2: Top 50 up-regulated genes in CIA SFs cultured in 2D.

| ID | log2FoldChange | foldChange | log10padj |
|---|---|---|---|
| Cdh4 | 4.172 | 18.020 | 34.632 |
| Cnmd | 3.218 | 9.303 | 75.540 |
| Chdh | 3.103 | 8.595 | 10.571 |
| Hsd11b2 | 2.809 | 7.007 | 43.675 |
| Hnf1b | 2.615 | 6.126 | 22.009 |
| Rgs5 | 2.607 | 6.093 | 93.313 |
| Cybrd1 | 2.563 | 5.911 | 60.321 |
| Cldn19 | 2.489 | 5.615 | 51.337 |
| Slc25a31 | 2.337 | 5.054 | 6.440 |
| Chrdl2 | 2.322 | 5.001 | 31.685 |
| Ces1f | 2.317 | 4.983 | 7.584 |
| Cxcl13 | 2.308 | 4.952 | 14.164 |
| Slc15a2 | 2.244 | 4.738 | 11.390 |
| Chl1 | 2.242 | 4.730 | 222.390 |
| Kcnip1 | 2.144 | 4.420 | 5.463 |
| Inmt | 2.083 | 4.236 | 21.077 |
| Mycn | 2.062 | 4.176 | 7.239 |
| Wnt10a | 2.047 | 4.133 | 23.225 |
| Kcnq4 | 1.997 | 3.991 | 7.006 |
| Pax8 | 1.933 | 3.818 | 80.990 |
| Crygs | 1.891 | 3.709 | 7.402 |
| Met | 1.870 | 3.656 | 92.430 |
| Col4a6 | 1.842 | 3.584 | 10.291 |
| Syndig1 | 1.785 | 3.447 | 44.148 |
| Syndig1 | 1.785 | 3.447 | 44.148 |
| Neto2 | 1.765 | 3.398 | 122.911 |
| Gjb3 | 1.765 | 3.398 | 38.109 |
| G0s2 | 1.764 | 3.396 | 13.440 |
| Cldn1 | 1.726 | 3.308 | 62.258 |
| Slc36a2 | 1.724 | 3.303 | 11.724 |
| Serpina3c | 1.721 | 3.297 | 12.927 |
| C1ql1 | 1.690 | 3.226 | 10.856 |
| Car3 | 1.592 | 3.014 | 19.758 |
| Shroom2 | 1.585 | 3.000 | 15.246 |
| Rbfox1 | 1.570 | 2.969 | 14.339 |
| Fmo1 | 1.564 | 2.957 | 21.882 |
| Plin1 | 1.562 | 2.953 | 14.152 |
| Clec3b | 1.559 | 2.947 | 13.578 |
| Cd24a | 1.548 | 2.925 | 47.913 |
| Gpr88 | 1.533 | 2.895 | 22.738 |
| Plac8 | 1.533 | 2.893 | 9.515 |
| Itga3 | 1.502 | 2.833 | 68.722 |
| Ccl11 | 1.499 | 2.825 | 6.344 |
| Rgs4 | 1.484 | 2.798 | 40.755 |
| Vat1l | 1.483 | 2.794 | 14.603 |
| Scin | 1.477 | 2.784 | 4.215 |
| Gm8113 | 1.477 | 2.784 | 6.275 |
| Mmp3 | 1.470 | 2.771 | 113.900 |

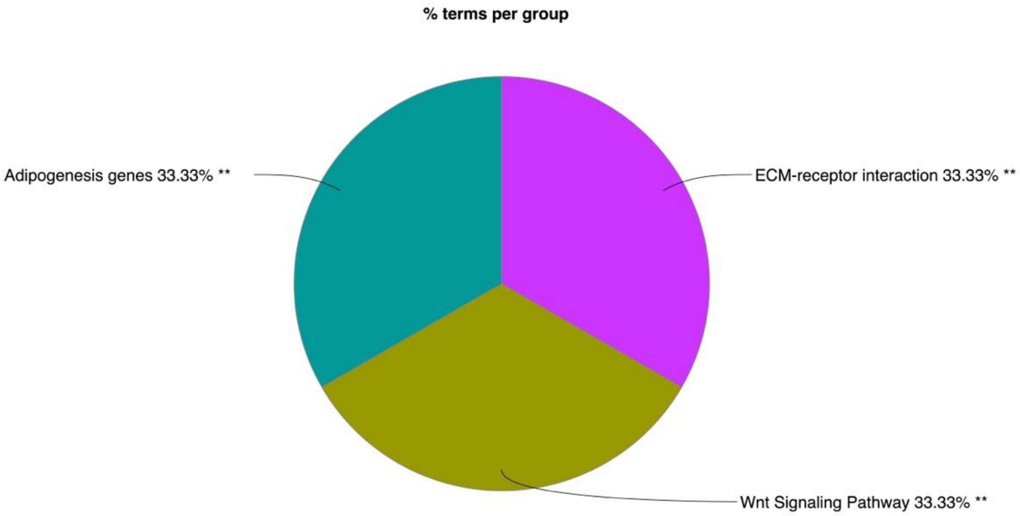

Supplementary Table 3: Top 50 up-regulated genes in CIA SFs cultured in scaffolds.

| ID | log2FoldChange | foldChange | log10padj |
|---|---|---|---|
| Csf3 | 6.828 | 113.622 | 4.380 |
| Nos2 | 4.732 | 26.575 | 3.467 |
| Il6 | 4.178 | 18.098 | 3.071 |
| 2010005H15Rik | 3.945 | 15.403 | 2.655 |
| Cxcl11 | 3.831 | 14.228 | 3.039 |
| Nrsn1 | 3.688 | 12.890 | 11.414 |
| Cxcl3 | 3.664 | 12.678 | 2.586 |
| Sfrp4 | 3.433 | 10.800 | 13.756 |
| U90926 | 3.305 | 9.882 | 12.726 |
| Mycn | 3.229 | 9.377 | 16.726 |
| Kcnip1 | 3.135 | 8.787 | 7.124 |
| Acod1 | 3.097 | 8.554 | 2.059 |
| Gjb3 | 2.896 | 7.442 | 30.011 |
| Mxd3 | 2.799 | 6.959 | 7.584 |
| Xkr5 | 2.798 | 6.956 | 10.103 |
| Cpm | 2.760 | 6.772 | 3.266 |
| Chrdl2 | 2.655 | 6.298 | 4.558 |
| Cdc25c | 2.649 | 6.273 | 7.744 |
| Lockd | 2.647 | 6.262 | 3.052 |
| Ckap2 | 2.644 | 6.252 | 3.822 |
| B4galnt3 | 2.622 | 6.157 | 17.838 |
| Cdc20 | 2.605 | 6.082 | 5.528 |
| Cxcl1 | 2.544 | 5.834 | 3.942 |
| Tinagl1 | 2.541 | 5.820 | 24.157 |

| ID | log2FoldChange | foldChange | log10padj |
|---|---|---|---|
| Cenpf | 2.537 | 5.805 | 3.563 |
| Kif20a | 2.522 | 5.745 | 4.315 |
| Ppl | 2.505 | 5.678 | 4.784 |
| Fam83d | 2.484 | 5.596 | 4.061 |
| Cdhr1 | 2.482 | 5.586 | 14.418 |
| Aspm | 2.476 | 5.565 | 4.054 |
| Kcnf1 | 2.472 | 5.550 | 14.391 |
| Ube2c | 2.472 | 5.548 | 12.990 |
| Anln | 2.461 | 5.508 | 5.044 |
| Ccnb1 | 2.435 | 5.409 | 12.726 |
| Prc1 | 2.399 | 5.276 | 4.443 |
| Apol9b | 2.398 | 5.270 | 7.425 |
| Styk1 | 2.392 | 5.250 | 8.029 |
| Troap | 2.391 | 5.245 | 7.195 |
| Gata2 | 2.388 | 5.235 | 41.006 |
| Hmmr | 2.378 | 5.197 | 4.266 |
| Ccdc18 | 2.370 | 5.168 | 4.579 |
| Cdh22 | 2.368 | 5.163 | 2.093 |
| Cdh26 | 2.364 | 5.146 | 2.973 |
| Chl1 | 2.357 | 5.123 | 68.780 |
| Ckap2l | 2.356 | 5.121 | 4.042 |
| Cd24a | 2.350 | 5.099 | 54.319 |
| Kif2c | 2.345 | 5.080 | 4.983 |
| Plk1 | 2.333 | 5.037 | 5.677 |

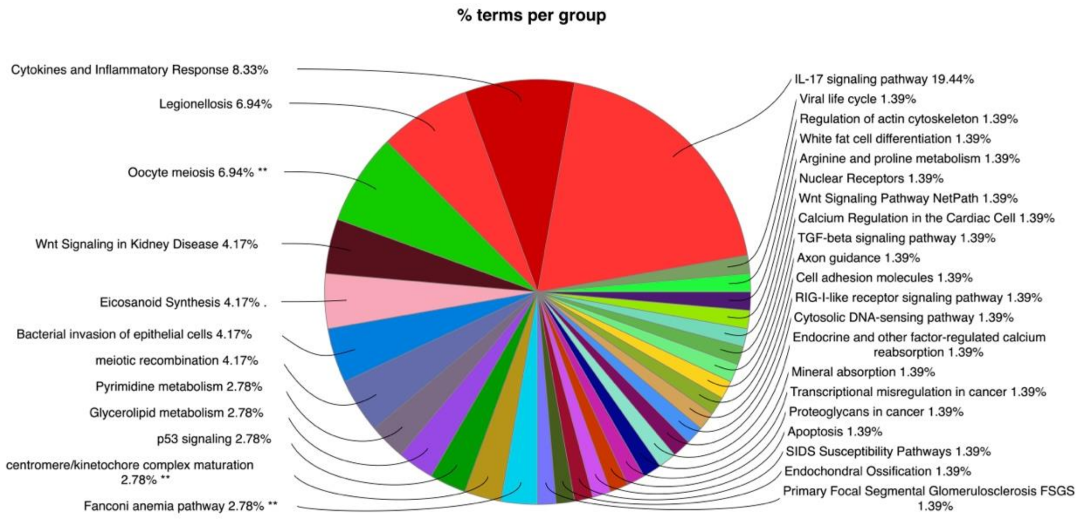

Supplementary Table 4: Top 50 up-regulated genes in CIA SFs cultured in hydrogels

| ID | log2FoldChange | foldChange | log10padj | ID | log2FoldChange | foldChange | log10padj |
|---|---|---|---|---|---|---|---|
| Spon2 | 4.228 | 18.735 | 5.229 | C4b | 2.297 | 4.916 | 18.775 |
| Sfrp4 | 3.877 | 14.688 | 11.138 | Sfrp2 | 2.199 | 4.590 | 2.089 |
| Avpr1a | 3.792 | 13.855 | 2.402 | Htra4 | 2.189 | 4.561 | 2.397 |
| Aspn | 3.572 | 11.893 | 3.356 | Olfml3 | 2.176 | 4.518 | 3.760 |
| Itm2a | 3.154 | 8.904 | 10.015 | Srpx | 2.167 | 4.492 | 4.708 |
| Cyp2f2 | 3.011 | 8.063 | 3.247 | Ndufa4l2 | 2.164 | 4.481 | 4.592 |
| Cthrc1 | 2.978 | 7.877 | 4.678 | Tmem100 | 2.138 | 4.402 | 3.703 |
| Mfap5 | 2.937 | 7.660 | 4.725 | Mgarp | 2.126 | 4.365 | 3.655 |
| Gm32219 | 2.931 | 7.629 | 5.602 | Scara5 | 2.057 | 4.161 | 7.619 |
| Clec11a | 2.922 | 7.578 | 3.401 | Islr | 2.050 | 4.140 | 3.374 |
| Tnfsf11 | 2.921 | 7.573 | 10.982 | Kng1 | 2.041 | 4.117 | 3.398 |
| Htra1 | 2.819 | 7.054 | 6.789 | Col1a2 | 2.041 | 4.115 | 4.095 |
| 3632451O06Rik | 2.799 | 6.958 | 2.842 | F13a1 | 2.031 | 4.088 | 6.174 |
| Col3a1 | 2.768 | 6.810 | 6.983 | Matn2 | 2.031 | 4.086 | 3.316 |
| Fxyd1 | 2.768 | 6.810 | 6.466 | Cox6b2 | 1.946 | 3.854 | 4.263 |
| Tnfaip6 | 2.634 | 6.209 | 10.775 | Adamts2 | 1.924 | 3.795 | 2.077 |
| Thy1 | 2.630 | 6.189 | 5.434 | Timp3 | 1.924 | 3.794 | 4.521 |
| Wisp2 | 2.625 | 6.168 | 6.496 | Egr2 | 1.911 | 3.760 | 2.984 |
| Tgfbi | 2.595 | 6.041 | 11.760 | Tshz2 | 1.892 | 3.711 | 3.116 |
| Arg1 | 2.584 | 5.995 | 4.579 | Rbp4 | 1.849 | 3.601 | 3.147 |
| Rarres2 | 2.430 | 5.390 | 2.658 | Pdlim2 | 1.835 | 3.567 | 3.532 |
| Tnfrsf11b | 2.391 | 5.244 | 4.707 | Mfap2 | 1.801 | 3.485 | 2.232 |
| Lrrc17 | 2.372 | 5.175 | 4.816 | Fbln1 | 1.780 | 3.435 | 4.286 |
| Col12a1 | 2.314 | 4.971 | 5.810 | Bgn | 1.779 | 3.433 | 6.993 |

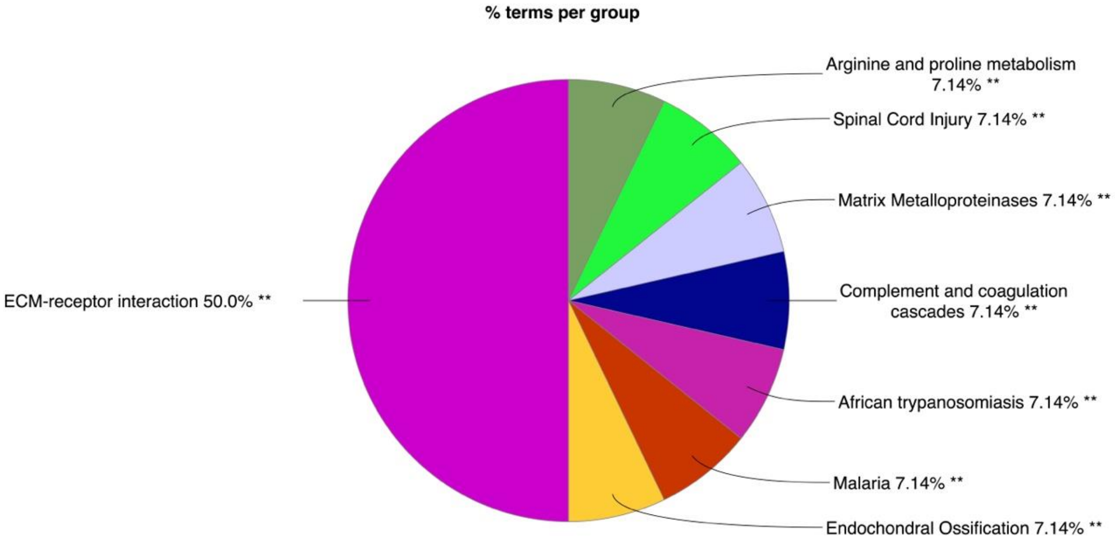

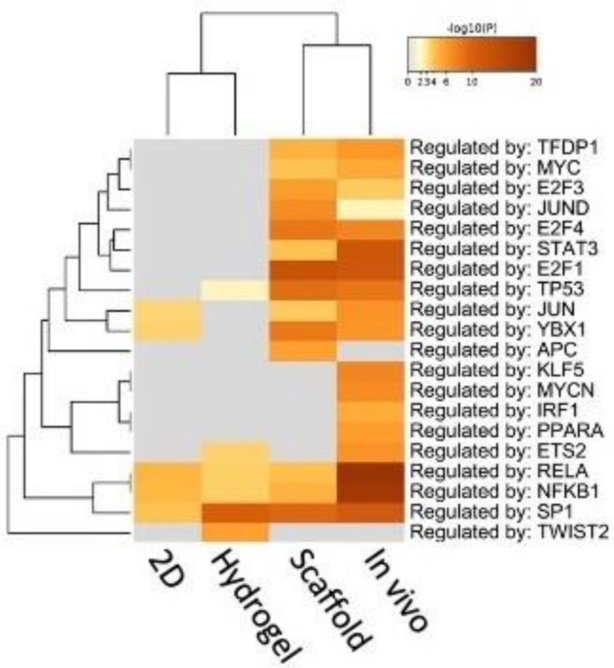


**Supplementary Figure 1. Comparative meta-analysis across in vivo, 2D and 3D cultured arthritic SFs RNA-Seq datasets**. Differentially expressed genes in CIA SFs compared to healthy controls in not cultured cells and cells cultured in 2D, 3D rigid scaffolds and FNPEG hydrogels were compared. TRRUST (Transcriptional Regulatory Relationships Unraveled by Sentence-based Text mining, https://www.grnpedia.org/trrust/) was used to identify activation of Transcription Factor-specific pathways in CIA SFs based on literature curation. The heatmap cells are colored by their p-values, white cells indicate the lack of enrichment for that term in the corresponding gene list. All results were generated with the bioinformatic tool Metascape.

### Supplementary Table 5: Top 100 up-regulated genes in IL-1-stimulated SFs

#### 2D cultured

| ID | log2FoldChange | foldChange | log10padj |
|---|---|---|---|
| Csf3 | 8.098 | 273.965 | 307.291 |
| Nos2 | 6.920 | 121.067 | 75.681 |
| Trdc | 6.223 | 74.698 | 41.906 |
| Csf2 | 6.122 | 69.629 | 36.004 |
| Fgf23 | 5.767 | 54.472 | 56.296 |
| Ccl20 | 5.489 | 44.915 | Inf |
| Cxcl1 | 5.297 | 39.303 | Inf |
| Il6 | 5.256 | 38.221 | Inf |
| Cxcl5 | 5.149 | 35.482 | Inf |
| Cxcl3 | 5.128 | 34.968 | Inf |
| Calcr | 5.066 | 33.509 | 28.150 |
| Gm16685 | 5.048 | 33.079 | 132.292 |
| Ccl5 | 4.701 | 26.009 | 256.968 |
| Mmp10 | 4.494 | 22.535 | 45.848 |
| Olr1 | 4.197 | 18.347 | 141.425 |
| Cxcl2 | 4.110 | 17.272 | 296.755 |
| Lif | 3.970 | 15.667 | Inf |
| Kcnk10 | 3.648 | 12.532 | 163.878 |
| Tnfsf15 | 3.428 | 10.766 | 238.502 |
| U90926 | 3.320 | 9.988 | 63.107 |
| Tac1 | 3.320 | 9.985 | 23.402 |
| Cxcl11 | 3.214 | 9.277 | 2.988 |
| Pdzd2 | 3.163 | 8.954 | 78.726 |
| Arg1 | 3.148 | 8.867 | 28.383 |
| Tmem132e | 3.144 | 8.838 | 56.903 |
| Serpinb2 | 3.135 | 8.788 | 231.701 |
| A730049H05Rik | 3.068 | 8.386 | 15.833 |
| Cxcl9 | 3.065 | 8.368 | 36.858 |
| Il1b | 3.024 | 8.133 | 248.739 |
| Ptgs2 | 3.017 | 8.095 | 309.342 |
| Dapk2 | 3.017 | 8.093 | 31.008 |
| Ccl2 | 3.008 | 8.044 | Inf |
| Has1 | 2.987 | 7.929 | 99.677 |
| Ereg | 2.954 | 7.752 | 221.498 |
| Fam167a | 2.915 | 7.544 | 69.417 |
| Syt7 | 2.902 | 7.475 | 28.332 |
| Steap4 | 2.845 | 7.183 | 192.197 |
| Procr | 2.776 | 6.851 | 170.434 |
| Slc1a2 | 2.747 | 6.713 | 20.671 |
| Acod1 | 2.726 | 6.616 | 3.608 |
| Selp | 2.707 | 6.532 | 19.237 |
| Zc3h12a | 2.703 | 6.511 | 299.594 |
| Upp1 | 2.634 | 6.207 | 52.149 |
| Mt2 | 2.578 | 5.972 | 186.529 |
| Tslp | 2.572 | 5.946 | 90.612 |
| Mmp3 | 2.565 | 5.916 | 279.612 |
| Il11 | 2.559 | 5.893 | 154.398 |
| Flt1 | 2.554 | 5.874 | 73.390 |
| Slc7a2 | 2.553 | 5.869 | Inf |
| Mmp13 | 2.536 | 5.798 | 176.882 |
| Kcnq5 | 2.532 | 5.783 | 38.681 |
| Camp | 2.530 | 5.777 | 22.359 |
| Ngf | 2.516 | 5.719 | 101.491 |
| Gem | 2.446 | 5.449 | 127.991 |
| Slc36a2 | 2.443 | 5.437 | 33.333 |
| Ccl11 | 2.430 | 5.391 | 26.475 |
| Rnd1 | 2.421 | 5.354 | 98.151 |
| Ier3 | 2.392 | 5.249 | 191.844 |
| Spink6 | 2.302 | 4.930 | 33.557 |
| Bdkrb1 | 2.298 | 4.916 | 91.716 |
| Cyp7b1 | 2.297 | 4.914 | 174.234 |
| Fmo1 | 2.263 | 4.801 | 67.172 |
| Slc7a11 | 2.261 | 4.794 | 118.639 |
| Lamb3 | 2.240 | 4.724 | 19.231 |
| Adora2b | 2.239 | 4.721 | 178.088 |
| Sod2 | 2.231 | 4.696 | 229.114 |
| Acpp | 2.189 | 4.559 | 31.490 |
| Adamts4 | 2.154 | 4.450 | 121.471 |
| Ifi205 | 2.148 | 4.433 | 33.475 |
| Egln3 | 2.139 | 4.405 | 65.712 |
| Ptges | 2.118 | 4.341 | 202.534 |
| Tnfaip3 | 2.118 | 4.340 | 179.182 |
| Dbx2 | 2.090 | 4.258 | 24.334 |
| Bdkrb2 | 2.055 | 4.155 | 79.685 |
| Abtb2 | 2.039 | 4.110 | 65.593 |
| Gm13889 | 2.038 | 4.107 | 17.753 |
| Spink5 | 2.035 | 4.098 | 7.062 |
| Artn | 2.020 | 4.055 | 32.379 |
| Gpr84 | 2.019 | 4.053 | 121.345 |
| Hs3st1 | 1.956 | 3.879 | 28.750 |
| Gbp6 | 1.955 | 3.878 | 9.844 |
| Penk | 1.946 | 3.854 | 181.970 |
| Nfkbiz | 1.926 | 3.801 | 101.564 |
| Ccl7 | 1.918 | 3.779 | 151.317 |
| Garem2 | 1.903 | 3.740 | 49.424 |
| Col23a1 | 1.889 | 3.704 | 95.614 |
| Lcn2 | 1.881 | 3.682 | 92.165 |
| Cd40 | 1.879 | 3.678 | 33.795 |
| Tnfsf11 | 1.864 | 3.641 | 63.610 |
| Gbp5 | 1.835 | 3.568 | 31.111 |
| Vcam1 | 1.832 | 3.560 | 202.568 |
| Nr4a2 | 1.828 | 3.549 | 111.269 |
| Arg2 | 1.824 | 3.540 | 17.078 |
| Gm26809 | 1.816 | 3.521 | 48.344 |
| Saa3 | 1.793 | 3.465 | 97.299 |
| Cpm | 1.758 | 3.383 | 19.498 |
| Eva1c | 1.745 | 3.351 | 10.786 |
| Cd69 | 1.741 | 3.342 | 14.998 |
| Marco | 1.730 | 3.317 | 13.656 |

#### Rigid scaffolds

| ID | log2FoldChange | foldChange | log10padj |
|---|---|---|---|
| Csf3 | 11.898 | 3816.370 | Inf |
| Nos2 | 9.790 | 885.478 | 111.765 |
| Csf2 | 9.257 | 611.873 | 34.511 |
| Il6 | 9.047 | 528.977 | Inf |
| Calcr | 8.600 | 387.989 | 48.512 |
| Cxcl3 | 8.545 | 373.498 | 296.639 |
| Rab44 | 7.795 | 222.078 | 50.058 |
| Ccl20 | 7.125 | 139.605 | 298.504 |
| Serpinb2 | 7.104 | 137.588 | 30.016 |
| Selp | 7.102 | 137.399 | 56.763 |
| Arg1 | 6.888 | 118.436 | 16.431 |
| Lif | 6.830 | 113.787 | 104.102 |
| Cxcl1 | 6.626 | 98.770 | 132.153 |
| Cilp2 | 6.520 | 91.782 | 75.673 |
| Il1b | 6.474 | 88.922 | 178.255 |
| Fgf23 | 6.103 | 68.757 | 65.661 |
| Saa3 | 6.027 | 65.208 | 32.396 |
| Has1 | 5.968 | 62.607 | 20.414 |
| Kcnk10 | 5.884 | 59.074 | 5.654 |
| Cxcl2 | 5.780 | 54.963 | 92.064 |
| Cxcl11 | 5.742 | 53.522 | 19.586 |
| Ptgs2 | 5.631 | 49.559 | 72.163 |
| Crispld2 | 5.415 | 42.668 | 118.926 |
| Tnfsf15 | 5.378 | 41.583 | 89.099 |
| Hpx | 5.297 | 39.328 | 213.738 |
| Gm16685 | 5.289 | 39.103 | 150.921 |
| Il11 | 5.275 | 38.719 | 37.362 |
| Rnd1 | 5.134 | 35.107 | Inf |
| 2010005H15Rik | 5.084 | 33.918 | 27.239 |
| Cxcl5 | 5.080 | 33.822 | Inf |
| Acod1 | 5.072 | 33.630 | 6.286 |
| Bdkrb1 | 5.004 | 32.086 | 155.689 |
| Cytip | 4.987 | 31.708 | Inf |
| Cdh22 | 4.957 | 31.060 | 30.709 |
| Cdh1 | 4.938 | 30.645 | 82.655 |
| Ccl5 | 4.924 | 30.355 | Inf |
| Gpr35 | 4.822 | 28.289 | 135.425 |
| Trem1 | 4.799 | 27.845 | 170.604 |
| Tmem132e | 4.717 | 26.305 | 84.305 |
| U90926 | 4.677 | 25.573 | 69.854 |
| Cpm | 4.604 | 24.317 | 17.361 |
| Ereg | 4.536 | 23.202 | 33.172 |
| Adora2b | 4.410 | 21.261 | Inf |
| Kcnh1 | 4.343 | 20.290 | 23.936 |
| Inhba | 4.149 | 17.742 | Inf |
| Ccl2 | 4.146 | 17.704 | Inf |
| Bdkrb2 | 4.115 | 17.328 | 121.901 |
| Egln3 | 4.103 | 17.187 | 122.237 |
| Dbx2 | 4.078 | 16.885 | 60.220 |
| Mt2 | 4.022 | 16.251 | Inf |
| Ctla2a | 4.010 | 16.107 | 318.110 |
| Cxcl9 | 3.997 | 15.963 | 6.807 |
| Ifi205 | 3.972 | 15.694 | 113.757 |
| Ret | 3.958 | 15.542 | 194.630 |
| Osbp2 | 3.928 | 15.220 | 56.055 |
| C430002N11Rik | 3.839 | 14.309 | 17.317 |
| Cxcl10 | 3.799 | 13.920 | 8.980 |
| Tgfbi | 3.785 | 13.780 | Inf |
| Ngf | 3.657 | 12.618 | 136.554 |
| Nr4a2 | 3.643 | 12.493 | 244.940 |
| Lcn2 | 3.589 | 12.033 | Inf |
| Arg2 | 3.582 | 11.978 | 20.529 |
| Nr4a3 | 3.577 | 11.932 | 63.802 |
| Ier3 | 3.531 | 11.562 | 80.516 |
| Gpr84 | 3.525 | 11.515 | 217.058 |
| Il4ra | 3.509 | 11.388 | Inf |
| F2rl2 | 3.485 | 11.196 | 18.214 |
| Kcnq5 | 3.485 | 11.194 | 55.653 |
| Ccl12 | 3.485 | 11.193 | 66.723 |
| Gfpt2 | 3.484 | 11.190 | Inf |
| Cass4 | 3.448 | 10.912 | 52.776 |
| Garem2 | 3.432 | 10.790 | 74.422 |
| Acat3 | 3.429 | 10.767 | 18.230 |
| Elovl7 | 3.427 | 10.756 | 50.491 |
| Adora3 | 3.413 | 10.654 | 21.652 |
| Lamb3 | 3.403 | 10.575 | 20.184 |
| Bmpr1b | 3.362 | 10.282 | 25.226 |
| Dusp5 | 3.319 | 9.978 | 293.123 |
| Pde4b | 3.302 | 9.865 | 274.437 |
| Zc3h12a | 3.299 | 9.840 | Inf |
| Socs3 | 3.295 | 9.814 | 315.718 |
| Nr4a1 | 3.264 | 9.609 | 48.690 |
| Upp1 | 3.258 | 9.569 | 49.534 |
| Ifi209 | 3.248 | 9.497 | 51.266 |
| Clec4e | 3.227 | 9.366 | 147.066 |
| Slc16a3 | 3.220 | 9.318 | 35.639 |
| Kcnn3 | 3.219 | 9.313 | 31.739 |
| Crabp2 | 3.217 | 9.296 | 194.405 |
| Slco2a1 | 3.216 | 9.292 | 77.657 |
| Camp | 3.190 | 9.123 | 20.348 |
| Runx3 | 3.188 | 9.113 | 95.566 |
| Ccl7 | 3.187 | 9.105 | Inf |
| Tm4sf1 | 3.172 | 9.011 | Inf |
| Galnt18 | 3.166 | 8.974 | 157.599 |
| Tnfsf11 | 3.148 | 8.863 | 100.311 |
| Ak3l2-ps | 3.143 | 8.831 | 33.237 |
| Mmp10 | 3.135 | 8.785 | 72.858 |
| Nt5e | 3.124 | 8.717 | 57.659 |
| Flt1 | 3.091 | 8.519 | 73.616 |

#### Hydrogels

| ID | log2FoldChange | foldChange | log10padj |
|---|---|---|---|
| Gm37703 | 10.099 | 1096.564 | 2.111 |
| Olr1 | 9.608 | 780.501 | 3.495 |
| Gm47796 | 9.338 | 647.074 | 2.486 |
| Entpd1 | 9.193 | 585.412 | 2.072 |
| Ccl20 | 5.293 | 39.198 | 66.010 |
| Cxcl2 | 4.169 | 17.990 | 30.158 |
| Csf2 | 3.942 | 15.371 | 3.523 |
| Csf3 | 3.916 | 15.099 | 2.377 |
| Ptx3 | 3.364 | 10.293 | 16.405 |
| Neurl3 | 3.303 | 9.868 | 12.599 |
| U90926 | 3.216 | 9.292 | 3.565 |
| Ccl2 | 3.052 | 8.291 | 21.856 |
| Il1f6 | 2.770 | 6.820 | 14.506 |
| Tgm2 | 2.766 | 6.804 | 2.072 |
| Cxcl1 | 2.744 | 6.700 | 22.051 |
| Il6 | 2.704 | 6.518 | 6.754 |
| Serpinb2 | 2.656 | 6.302 | 4.731 |
| Mmp1b | 2.606 | 6.086 | 5.414 |
| Gpr84 | 2.593 | 6.033 | 2.555 |
| Ccl3 | 2.532 | 5.784 | 7.892 |
| Gm16685 | 2.516 | 5.720 | 2.425 |
| Fam84a | 2.495 | 5.636 | 5.307 |
| Nabp1 | 2.478 | 5.571 | 2.190 |
| Slfn2 | 2.424 | 5.367 | 4.025 |
| Ereg | 2.353 | 5.110 | 33.855 |
| Cxcl3 | 2.261 | 4.794 | 2.016 |
| Tnfaip3 | 2.189 | 4.560 | 5.468 |
| Stx11 | 2.179 | 4.527 | 5.517 |
| Kcnk10 | 2.153 | 4.447 | 3.986 |
| Ccl7 | 2.057 | 4.161 | 3.554 |
| Ch25h | 2.026 | 4.073 | 5.468 |
| Rgs16 | 2.021 | 4.060 | 4.663 |
| Lif | 1.961 | 3.892 | 5.176 |
| Twist2 | 1.849 | 3.602 | 3.488 |
| Ccl5 | 1.842 | 3.585 | 4.892 |
| Ptgs2 | 1.811 | 3.510 | 5.045 |
| Gbp2 | 1.791 | 3.461 | 6.583 |
| Itga2 | 1.781 | 3.436 | 2.840 |
| Plaur | 1.781 | 3.436 | 5.468 |
| Icam1 | 1.774 | 3.420 | 3.161 |
| Arg2 | 1.722 | 3.299 | 4.284 |
| Gch1 | 1.695 | 3.237 | 3.038 |
| Cxcl5 | 1.670 | 3.183 | 4.348 |
| Rel | 1.651 | 3.140 | 4.896 |
| Upp1 | 1.650 | 3.138 | 2.343 |
| Ehd1 | 1.641 | 3.118 | 4.575 |
| Nfkb1 | 1.619 | 3.072 | 3.036 |
| Plat | 1.595 | 3.021 | 3.380 |
| Trdc | 1.585 | 3.000 | 3.428 |
| Trdj1 | 1.577 | 2.984 | 2.147 |
| Bdkrb2 | 1.540 | 2.907 | 3.001 |
| Cxcl16 | 1.538 | 2.905 | 6.821 |
| Nfkbib | 1.526 | 2.879 | 2.111 |
| Ets1 | 1.489 | 2.807 | 3.843 |
| Il11 | 1.454 | 2.740 | 4.317 |
| Ier5 | 1.443 | 2.719 | 4.140 |
| Nfkbia | 1.440 | 2.713 | 6.830 |
| Il1rl1 | 1.406 | 2.650 | 5.908 |
| Flrt3 | 1.373 | 2.590 | 7.949 |
| Nrg1 | 1.365 | 2.575 | 3.166 |
| Slc7a2 | 1.353 | 2.555 | 5.468 |
| Mmp10 | 1.346 | 2.542 | 3.307 |
| Cyr61 | 1.327 | 2.509 | 2.377 |
| Nfkbiz | 1.292 | 2.449 | 5.524 |
| Plk2 | 1.291 | 2.448 | 4.289 |
| Vcam1 | 1.290 | 2.445 | 3.978 |
| Inhba | 1.279 | 2.426 | 4.207 |
| Slc7a11 | 1.244 | 2.369 | 7.280 |
| Odc1 | 1.240 | 2.361 | 2.124 |
| Plscr1 | 1.238 | 2.359 | 2.779 |
| Hbegf | 1.231 | 2.348 | 3.971 |
| Pde4b | 1.163 | 2.240 | 2.506 |
| Bcl10 | 1.152 | 2.223 | 2.586 |
| Rab8b | 1.141 | 2.205 | 3.103 |
| Glrx | 1.130 | 2.188 | 4.379 |
| Gfpt2 | 1.107 | 2.154 | 4.348 |
| Pdpn | 1.090 | 2.129 | 2.555 |
| Il1rap | 1.077 | 2.109 | 2.343 |
| Btg2 | 1.062 | 2.087 | 2.313 |
| Hnrnpab | 1.032 | 2.045 | 2.467 |
| Pip5k1a | 1.016 | 2.023 | 3.410 |
| Pla2g4a | 1.008 | 2.011 | 3.752 |
